# A periplasmic protein complex supports arabinofuranosyltransferase activity and mediates intrinsic drug resistance in *Mycobacterium tuberculosis*

**DOI:** 10.1101/2025.07.08.663683

**Authors:** Thaís Klevorn, Christopher Brown, Christine D. Hardy, Bonnie J. Cuthbert, Amanda Spencer, Adrián Jinich, Luming Chan, Shiva K. Angala, Jordan Manzer, Jessica Mendoza, Rodger de Miranda, Heather Kim, Dirk Schnappinger, Mary Jackson, Kyu Rhee, Celia W. Goulding, Sabine Ehrt

**Affiliations:** Department of Microbiology and Immunology, Weill Cornell Medical College, New York, New York, USA; Immunology and Microbial Pathogenesis Graduate Program, Weill Cornell Graduate School of Medical Sciences, Cornell University, New York, New York, USA; Department of Medicine, Weill Cornell Medical College, New York, New York, USA; Department of Molecular Biology & Biochemistry, University of California Irvine, Irvine, California, USA; Department of Chemistry & Biochemistry, Skaggs School of Pharmacy & Pharmaceutical Sciences, University of California San Diego, San Diego, California, USA; Mycobacteria Research Laboratories, Department of Microbiology, Immunology and Pathology, Colorado State University, Fort Collins, Colorado, USA; Department of Pharmaceutical Sciences, University of California Irvine, Irvine, California, USA

## Abstract

The intrinsic drug resistance of *Mycobacterium tuberculosis* (Mtb) is a major barrier to effective tuberculosis (TB) treatment, largely due to its complex, impermeable cell envelope. We identified a periplasmic protein complex comprising FecB and Rv3035 that is essential for maintaining envelope integrity and mediating intrinsic multidrug resistance in Mtb. FecB interacts with Rv3035, forming a stable heterodimer that associates with the cell envelope biosynthesis protein AftB. We report the structures of Rv3035 alone and in complex with FecB and identify critical residues for complex formation and function. Co-essentiality and genetic interaction analyses support a functional link between FecB, Rv3035 and AftB, an arabinofuranosyltransferase which synthesizes arabinogalactan and lipoarabinomannan. Loss of FecB or Rv3035 disrupted AftB-mediated arabinan synthesis, suggesting that these proteins support AftB’s enzymatic activity. Importantly, FecB is required for Mtb virulence in mice, underscoring its physiological relevance. These findings highlight FecB, Rv3035 and AftB as promising therapeutic targets.

## Introduction

Tuberculosis (TB) remains the world’s leading cause of death from a single infectious agent and is responsible for over 1 million deaths annually (*1*). Unlike most bacterial infections, successful TB treatment requires multiple antibiotics taken for several months, even in drug-susceptible cases. This prolonged therapy is in part due to the intrinsic drug resistance of *Mycobacterium tuberculosis* (Mtb), the causative agent of TB (*2*). A key contributor to this resistance is Mtb’s complex, multi-layered cell envelope, which acts as a protective barrier against antibiotics and host immune defenses (*3*). Despite its central role in Mtb’s pathogenicity and drug resistance, the molecular mechanisms governing the structure, assembly, and function of the mycobacterial cell envelope remain incompletely understood (*4*).

To identify novel targets that could potentiate existing TB therapies and shorten treatment duration, we previously performed a genome-wide chemical-genetic screen using transposon mutagenesis and sequencing (*5*). This identified *fecB* (Rv3044) as a gene whose disruption sensitized Mtb to multiple antibiotics with various mechanisms of action. These included rifampicin and ethambutol, which are part of the standard treatment regimen for drug-susceptible TB, and pretomanid, which is part of the BPaL (bedaquiline, pretomanid, linezolid) regimen used to treat drug-resistant TB (*5*). Functional studies of a Mtb *fecB* knockout strain (Δ*fecB*) revealed that *fecB* is essential for maintaining the impermeability of the Mtb cell envelope (*5*).

In Mtb, FecB is annotated as a periplasmic substrate-binding protein homologous to FecB from *Escherichia coli*, which mediates ferric citrate transport (*6*). However, Mtb lacks several canonical components of the ferric citrate transport system. Instead, Mtb FecB binds heme and the siderophore carboxymycobactin and interacts with components of the mycobacterial siderophore transport machinery (*7*). Despite these biochemical insights, the physiological role of FecB in Mtb is not fully defined. Here, we investigated how FecB contributes to Mtb’s intrinsic antibiotic resistance and uncovered a critical functional interaction with Rv3035, a previously uncharacterized periplasmic protein. Rv3035 not only interacts with FecB, but also with the arabinofuranosyltransferase AftB (Rv3805c), bridging these two proteins to form a larger complex that is important for cell envelope biogenesis.

Given the urgent need for shorter and more effective TB treatments, elucidating mechanisms that underlie Mtb’s intrinsic drug resistance is of high therapeutic relevance (*2*, *3*). By characterizing the function of FecB and Rv3035 in Mtb, this study advances our understanding of periplasmic proteins in mycobacteria and highlights these proteins as promising targets for combination therapies aimed at enhancing the efficacy of existing anti-TB drugs.

## Results

### MtbΔ*fecB* displays a transcriptional iron deprivation signature and accumulates siderophores

Transcriptome analysis can provide functional insights by identifying pathways that are differentially regulated in a mutant strain compared to a wild type (WT) strain. These changes may reflect the mutant strain’s need to adapt in the absence of the gene of interest (*8*). To gain insight into the function of FecB, we performed transcriptomic analysis by RNA sequencing of Mtb WT, Δ*fecB* and a complemented strain (Δ*fecB::fecB,* containing a chromosomally integrated plasmid expressing *fecB* from the *hsp60* promoter). We decided to compare the transcriptional profile of the strains in regular growth conditions with detergent (0.05% Tween80) and without detergent, because the presence of Tween80, although helpful to prevent excessive clumping of mycobacterial cultures, also strips the mycobacterial capsule and some surface exposed lipids (*9*), making it a less physiologically relevant condition.

Compared to WT and the complemented strain, Δ*fecB* significantly upregulated the *iniBAC* operon, which is known to be induced in response to cell wall stress (*10*), both in the presence and absence of detergent in the medium (Fig. 1A and data S1). Furthermore, in detergent-free conditions, 61% of the upregulated (22 of 36) and 13% of downregulated (3 of 23) differentially expressed genes (DEGs) were related to iron utilization. The upregulated DEGs included all 14 siderophore biosynthesis genes (*mbtA*-*mbtN*), the genes that encode the IrtA/B transporter for siderophore uptake (*irtA*, *irtB*) and the ESX-3 secretion system substrates (*esxG*, *esxH*), which are required for mycobactin-mediated iron acquisition. The downregulated DEGs included *bfrB*, encoding the iron storage protein bacterioferritin (Fig. 1, A and B). When grown in detergent-free medium, Δ*fecB* thus displays a transcriptional iron deprivation signature even though the medium contains a high concentration of iron. This iron deprivation signature is conditional, and it was not present when Tween80 was included in the medium (Fig. 1A). One potential explanation for this difference in transcriptional response is that Tween80’s stripping of the capsule and some outer membrane lipids might facilitate uptake of iron in Δ*fecB* during growth in regular medium.

**Figure 1.**
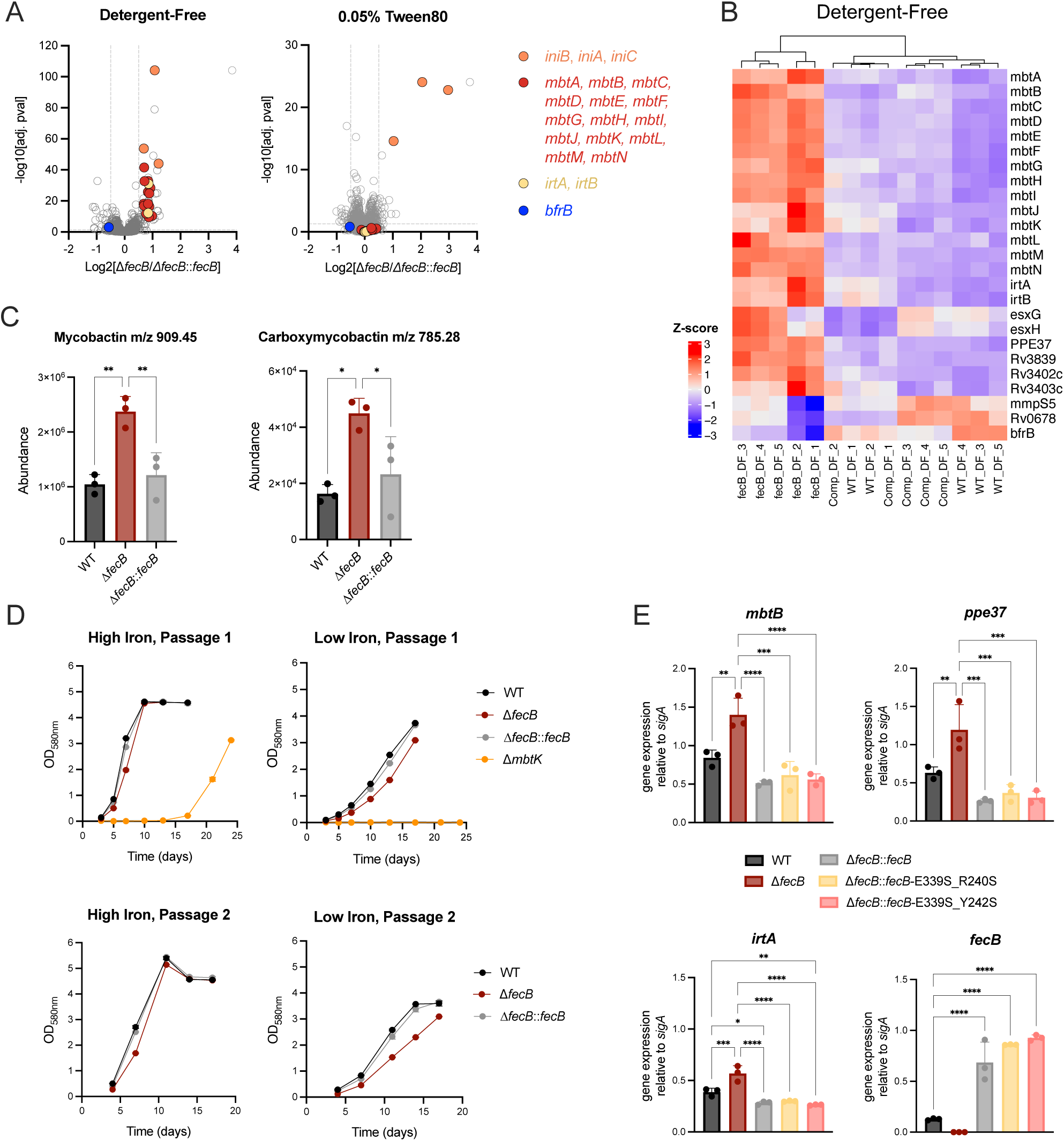
FecB plays a non-critical role in iron homeostasis in Mtb. (A) Volcano plots of transcriptome changes in Δ*fecB* compared to the complemented strain (Δ*fecB*::*fecB*) during log-phase growth in regular media without detergent or with 0.05% Tween80; highlighted genes are also differentially expressed in Δ*fecB* vs WT. (B) Heatmap of iron-related, differentially expressed genes (DEG) in strains grown in detergent-free media. DEGs were selected by padj ≤ 0.05 and |log2FC| ≥ 0.5; common hits in Δ*fecB* vs Δ*fecB*::*fecB* and Δ*fecB* vs WT are shown. (C) Relative abundance of representative mycobactin and carboxymycobactin species detected as [M+Fe- 2H]+ from lipid extracts analyzed by liquid chromatography-mass spectrometry (LC-MS). (D) Growth curves of iron pre-depleted strains in iron-chelated media with 160 µM (high iron) or 1 µM (low iron) ammonium ferric citrate (AFC). Cultures in low iron were grown until day 17 (Passage 1) and diluted to starting OD=0.025 in high/low iron media (Passage 2). Δ*mbtK* (mycobactin biosynthesis mutant) was used as a positive control that cannot grow in iron-limiting conditions. (E) Differential expression by reverse transcription quantitative PCR (RT qPCR) of representative iron-related genes (*mbtB*, *ppe37* and *irtA*) and *fecB* in strains grown in detergent-free media. Statistical significance in (C) and (E) was determined by one-way ANOVA and Tukey post-hoc test. Error bars: ±SD. \**P* < 0.05; \*\**P* < 0.01; \*\*\**P* < 0.001; \*\*\*\**P* <0.0001. N independent experiments = 2 with a total of five replicate cultures (A-B), 2 (C), 5 (D) and 1 (E) each performed with triplicate cultures.

Further supporting the transcriptomics data, the iron starvation signature in Δ*fecB* in detergent-free medium was accompanied by an accumulation of cell-associated siderophores as identified by mass-spectrometry analysis of lipid extracts (Fig. 1C). Taken together, these data suggest that Δ*fecB* may be less efficient in acquiring iron in detergent-free medium due to interrupted siderophore export through the periplasm, mycobacterial outer membrane and/or capsule.

### FecB is non-critical for mycobactin-mediated iron acquisition in Mtb

To further probe FecB’s potential role in iron acquisition, we analyzed the growth of Δ*fecB*, WT and Δ*fecB*::*fecB* in iron-limiting conditions (Fig. 1D). We included a siderophore mutant, MtbΔ*mbtK*, to ensure that the growth medium was iron limiting. After starving the strains for iron to deplete internal iron stores and then growing them in either high or low iron conditions (iron-chelated medium with 160 µM or 1 µM ammonium ferric citrate, respectively), we observed only a minor growth defect of Δ*fecB* compared to the control strains, both in the presence (Fig. 1D) or absence of detergent (fig. S1A). Upon passaging the strains growing in low iron into both high and low iron conditions, the growth defect was still subtle and much weaker than the strong growth defect of Δ*mbtK* (Fig. 1D).

FecB can bind to ferric- and apo-carboxymycobactin with high affinity *in vitro*, and this binding is disrupted by mutating key residues E339, R240 and Y242 (*7*). To test whether binding to siderophores is required for complementing Δ*fecB*’s iron-related phenotypes, we cloned and expressed FecB variants harboring the mutations E339S together with R240S or Y242S in Δ*fecB*. The FecB variants with significantly reduced binding affinity to ferric-carboxymycobactin (E339S_R240S and E339S_Y242S) fully reversed the iron deprivation transcriptional signature (examined by representative gene expression of *mbtB*, *ppe37* and *irtA*) in detergent-free medium (Fig. 1E) and the subtle growth defect in low iron (fig. S1B). This complementation was particularly remarkable given that the variant proteins were not clearly detectable in whole cell lysates via western blot (fig. S1C). However, we note that the FecB E339S_R240S and E339S_Y242S double mutants bind apo-carboxymycobactin with similar (or slightly higher) affinity than WT FecB (fig. S1D). Thus, we cannot rule out that binding apo-siderophores is an important function of FecB.

Taken together, these results suggest that FecB plays a role in iron acquisition in Mtb, however, the subtle growth defect of Δ*fecB* in iron-limiting conditions points to a condition-dependent non-critical function that is likely compensated for by other proteins required for iron acquisition.

### FecB’s role in intrinsic multidrug resistance is distinct from its activity in iron homeostasis

FecB is critical for Mtb’s cell envelope impermeability and resistance to multiple antibiotics (*5*). To test whether FecB’s activity in iron homeostasis is related to its activity in mediating cell envelope integrity and multidrug resistance in Mtb, we tested whether the FecB variants with reduced binding to ferric-carboxymycobactin could rescue the antibiotic hypersusceptibility of Δ*fecB*. Vancomycin and rifampicin were chosen for minimal inhibitory concentration (MIC) assays because Δ*fecB* is significantly more susceptible to these drugs compared to WT(*5*). Complementation of Δ*fecB* with the FecB double mutants (E339S_R240S and E339S_Y242S) fully rescued Δ*fecB*’s hypersusceptibility to vancomycin and rifampicin (Fig. 2A) indicating that ferric-carboxymycobactin binding is not required for FecB’s function in mediating antibiotic resistance.

**Figure 2.**
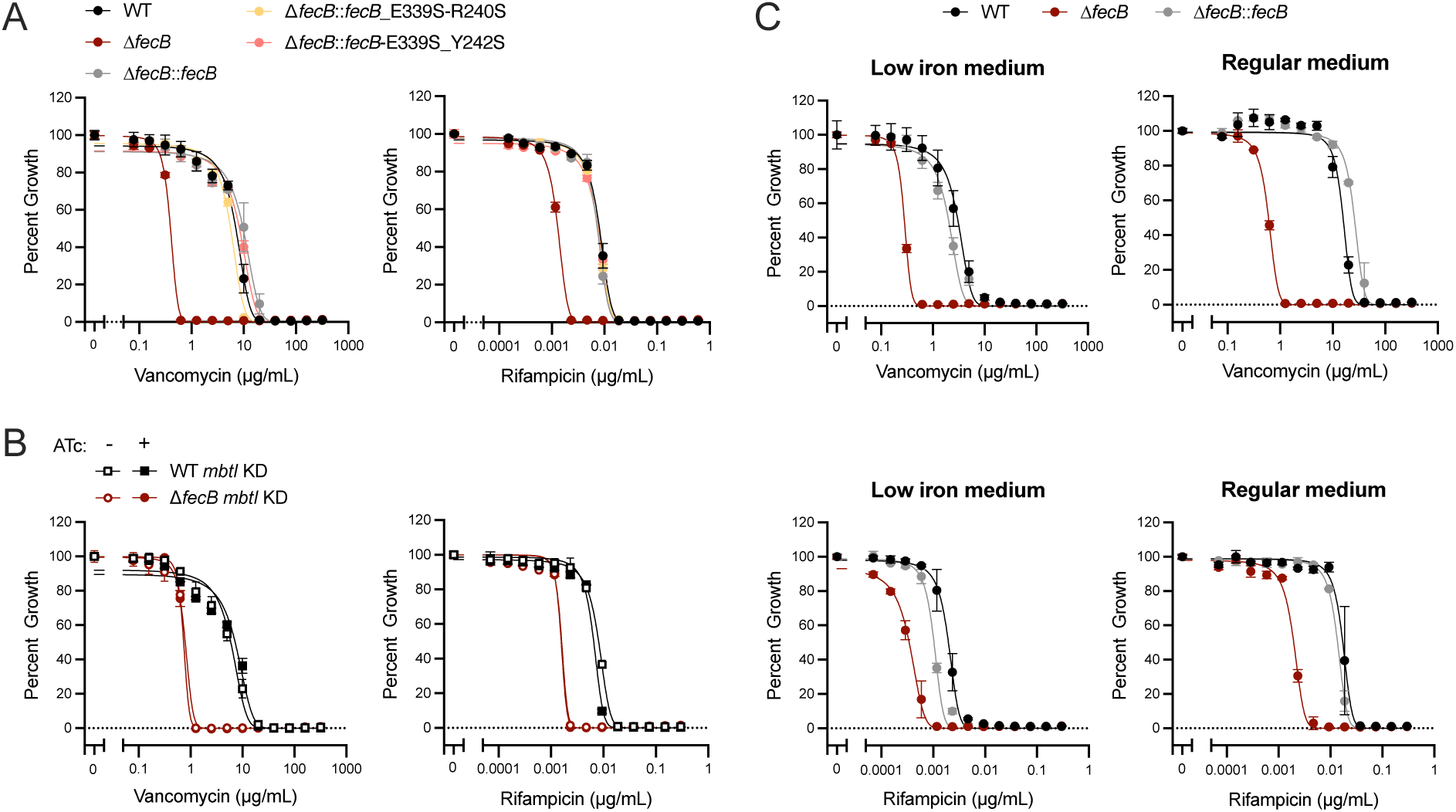
FecB’s roles in iron homeostasis and multidrug resistance are not functionally related. MIC assays for vancomycin and rifampicin, percent growth calculated from no drug control wells. (A) Δ*fecB* complemented with FecB variants with lower binding affinity to ferric carboxymycobactin (Δ*fecB*::*fecB*-E339S_R240S and Δ*fecB*::*fecB*-E339S_Y242S) and control strains (day 14). (B) CRISPRi KD of *mbtI* in WT/Δ*fecB* in the absence or presence of ATc inducer (day 14). (C) WT, Δ*fecB* and Δ*fecB*::*fecB* strains in low iron (iron-chelated media with 1µM AFC) or regular media (day 21). Error bars: ±SD. N independent experiments = 2 (A, C), 1 (B), each performed with triplicate cultures.

To explore whether siderophore accumulation in Δ*fecB* (Fig. 1C) leads to antibiotic hypersusceptibility, we interrupted siderophore biosynthesis via CRISPR interference (CRISPRi) knockdown (KD) of the mycobactin biosynthesis gene *mbtI* in Δ*fecB*. If siderophore accumulation is directly related to increased antibiotic susceptibility in Δ*fecB*, reducing siderophore production by knocking down *mbtI* should relieve this phenotype. We validated the KD of *mbtI* in WT and Δ*fecB* backgrounds by growing the strains in iron-limiting conditions, in the presence or absence of anhydrotetracycline (ATc) to activate CRISPRi (fig. S2A). In the presence of ATc, the *mbtI* KD strains failed to grow with ammonium ferric citrate concentrations lower than 1 µM, similarly to Δ*mbtK*, indicating that siderophore production is significantly impaired through silencing of *mbtI*. We next measured the MICs of vancomycin and rifampicin against the *mbtI* KD strain in the presence or absence of ATc (Fig. 2B). Silencing *mbtI* with ATc did not alter the MICs, indicating that siderophore accumulation is likely not the cause of antibiotic hypersusceptibility in Δ*fecB*.

To further test whether siderophores are involved in the antibiotic hypersusceptibility of Δ*fecB*, we tested whether induction of siderophore biosynthesis would enhance Δ*fecB*’s susceptibility to vancomycin and rifampicin. We performed MIC assays in low iron medium, where siderophore synthesis is induced, and in regular medium, where siderophore synthesis is suppressed (Fig. 2C and fig. S2B). Additionally, since an iron-starvation transcriptional signature, which includes the overexpression of siderophore biosynthesis genes, was observed in Δ*fecB* only in the absence of detergent (Fig. 1A), we also tested whether the mutant’s antibiotic hypersusceptibility increased in detergent-free medium compared to medium containing 0.05% Tween80 (fig. S2C). Contrary to what would be expected if siderophores promote antibiotic hypersusceptibility, the differences of the vancomycin and rifampicin MICs against Δ*fecB* compared to WT and Δ*fecB*::*fecB* decreased in low iron medium compared to regular iron rich medium (Fig. 2C). Similarly, a reduced shift of MICs was observed in detergent-free conditions (fig. S2C). These results do not support a direct involvement of siderophores in the antibiotic hypersusceptibility phenotype of Δ*fecB.* The reduced differences in antibiotic MICs might be due to slowed growth rates of the strains in low iron medium and increased clumping in detergent-free medium, which hinder accurate optical density measurements. In summary, these experiments dissociate FecB’s role in iron homeostasis from that in antibiotic resistance.

### FecB and Rv3035 interact in Mtb and likely function in the same cellular pathway

FecB is classified as a periplasmic substrate-binding protein that has been shown to interact with siderophore-mediated iron transport proteins (*7*). Since our data point to an alternative role for FecB in mediating intrinsic drug resistance in Mtb, we set out to identify additional physical interactors of FecB that could shed light on other pathways in which FecB might participate. Immunoprecipitation and mass spectrometry analysis of FLAG-tagged FecB was performed and analyzed in comparison to the immunoprecipitation of a WT lysate control to eliminate non-specific interactions.

The most highly enriched interactor in our dataset was the uncharacterized protein Rv3035 (Fig. 3A). Rv3035 is annotated as an essential hypothetical protein containing WD40 repeats that together are predicted to form an eight-bladed β-propeller structure. These domains can serve as scaffolds in protein-protein interactions and the formation of multiprotein complexes and play important roles in prokaryotes and eukaryotes (*11*). We confirmed binding of FecB to Rv3035 by co-immunoprecipitation of the tagged proteins (fig. S3) and then set out to identify Rv3035’s physical interactors by using Rv3035 as the bait in a second immunoprecipitation experiment. Satisfyingly, FecB was the most enriched protein in the immunoprecipitate of Rv3035-3xFLAG, compared to the WT lysate control (Fig. 3B). In addition, AftB and Rv0227c, which are the other two most enriched interacting partners of FecB, were also among the most enriched partners of Rv3035 (Fig. 3, A and B, and data S2).

**Figure 3.**
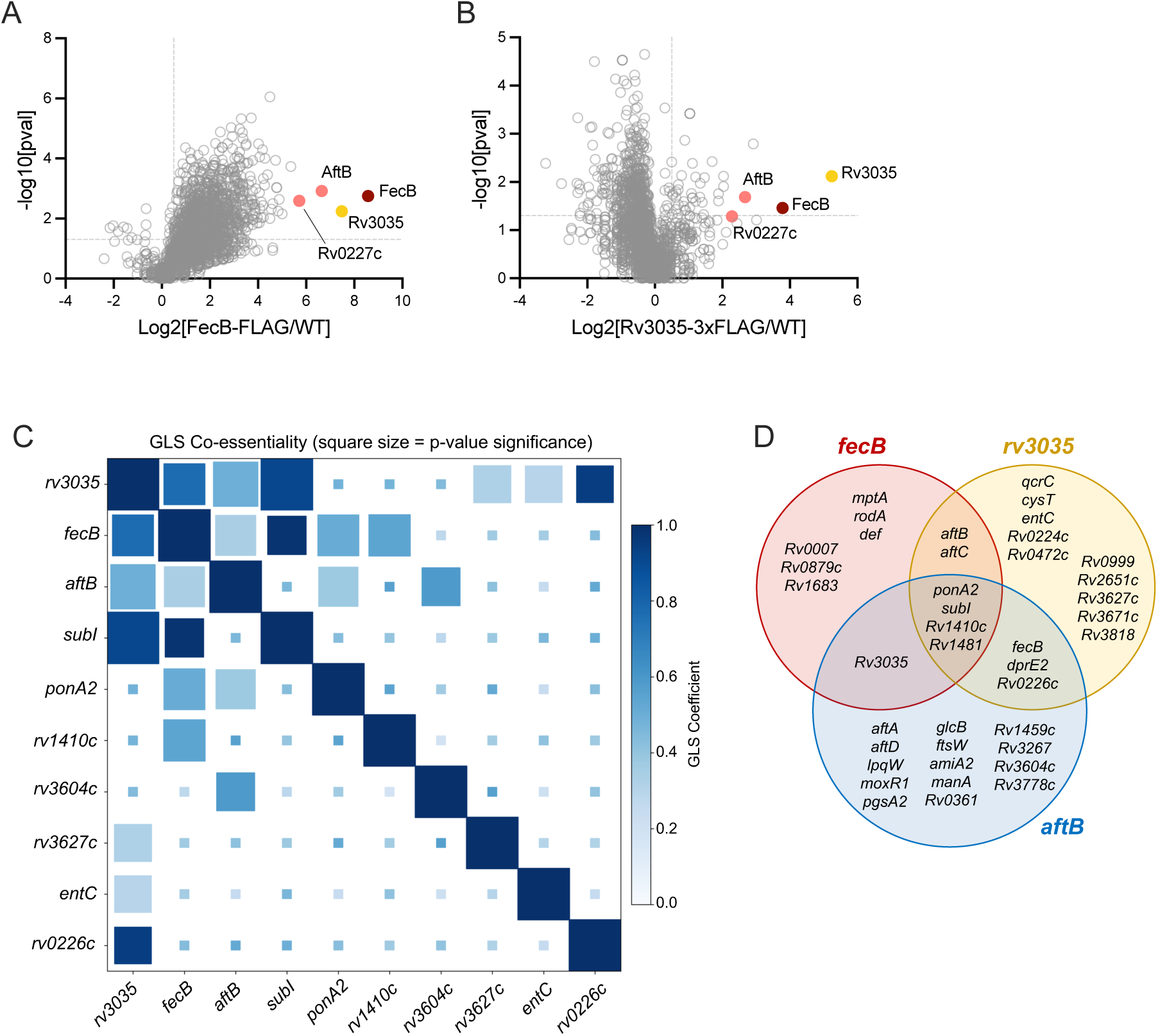
FecB and Rv3035 interact in Mtb and share essentiality profiles. Protein interaction profiling of (A) FecB and (B) Rv3035. FecB-FLAG and Rv3035-3xFLAG were immunoprecipitated from live Mtb and interacting proteins were identified by mass spectrometry. WT lysate was used a control for unspecific interactions. (C) Heatmap of co-essentiality Generalized Least Squares (GLS) coefficients for selected gene pairs. The genes shown were selected based on three “query genes”: *fecB*, *rv3035*, and *aftB*. We included all gene pairs involving at least one of these queries that met a significance threshold of FDR-adjusted p-value < 0.001. Each square’s color represents the GLS coefficient (i.e., the strength of co-essentiality), while size corresponds to statistical significance (−log₁₀ of the FDR-adjusted p-value). Diagonal elements represent self-comparisons (each gene’s log₂ fold change profile against itself); while biologically uninformative, they are displayed at full size as a visual benchmark for perfect correlation. Some gene-gene combinations did not pass the significance threshold and are shown with smaller squares. (D) Venn diagram showing genes that are co-essential with *fecB* (red circle), *rv3035* (yellow circle) and *aftB* (blue circle), and their overlap. N independent experiments = 1 (A-B), each performed with triplicate cultures.

To further inform our proteomics findings, we investigated the genetic interactions of *fecB* via co-essentiality/co-fitness analysis. This method allows the detection of gene pairs that share similar fitness profiles across forward genetic screens, and thus can help identify genes that encode proteins with similar or related function (*12*). Using a publicly available CRISPRi chemical-genetic interaction dataset which profiled gene essentiality across 80 drug-treatment and CRISPRi pre-depletion conditions (*13*), we performed co-essentiality analysis using generalized least squares (GLS) regression of *fecB*, *rv3035*, and the genes encoding their most enriched common protein interactors, *aftB* and *rv0227c*. While *rv0227c* did not share an essentiality profile with the others, *fecB*, *rv3035* and *aftB* were all hits in each other’s co-essential gene clusters (Fig. 3, C and D). This analysis identified 13 genes co-essential with *fecB*, 19 with *rv3035* and 22 with *aftB*, among which 4 were shared by all three (Fig. 3D and data S3).

### FecB forms a complex with Rv3035 *in vitro*

To confirm the direct binding of Rv3035 and FecB *in vitro*, we cloned His-tagged Rv3035 from *Mycobacterium thermoresistible* (Mth). Mth Rv3035 shares 67% sequence identity with Mtb Rv3035 (fig. S4A) and was used because full-length or N-terminal truncation constructs of His-tagged Mtb Rv3035 failed to yield soluble protein. Mth Rv3035 is predicted to have an N-terminal signal peptide by SignalP 6.0 (*14*) and to be a lipoprotein, where the N-terminal cysteine residue is predicted to form a thioether linkage to diacylglycerol (*15*, *16*). Here, we have designated this N-terminal cysteine as the first amino acid of the mature form of Rv3035 (fig. S4B). Our Mth Rv3035 protein construct begins at residue 2 of the predicted mature protein and was expressed with a cleavable, N-terminal His-tag. Mth Rv3035-His was purified to homogeneity and the His-tag cleaved, with size-exclusion chromatography (SEC) of the resulting protein suggesting that Mth Rv3035 is monomeric (fig. S5A).

Mth Rv3035-His was also co-expressed with untagged Mtb FecB and found to co-elute by Ni-affinity chromatography (fig. S5B). The Mth Rv3035 and Mtb FecB complex was stable during SEC and displayed a molecular weight of 79.1 kDa, consistent with a heterodimer of Rv3035 and FecB (fig. S5A). The affinity of Mth Rv3035 for Mtb FecB was quantified using isothermal titration calorimetry (ITC). This analysis revealed tight 1:1 binding of Mth Rv3035 and Mtb FecB, with a K_D_ of 27.5 ± 4.6 nM (Fig. 4A).

**Figure 4.**
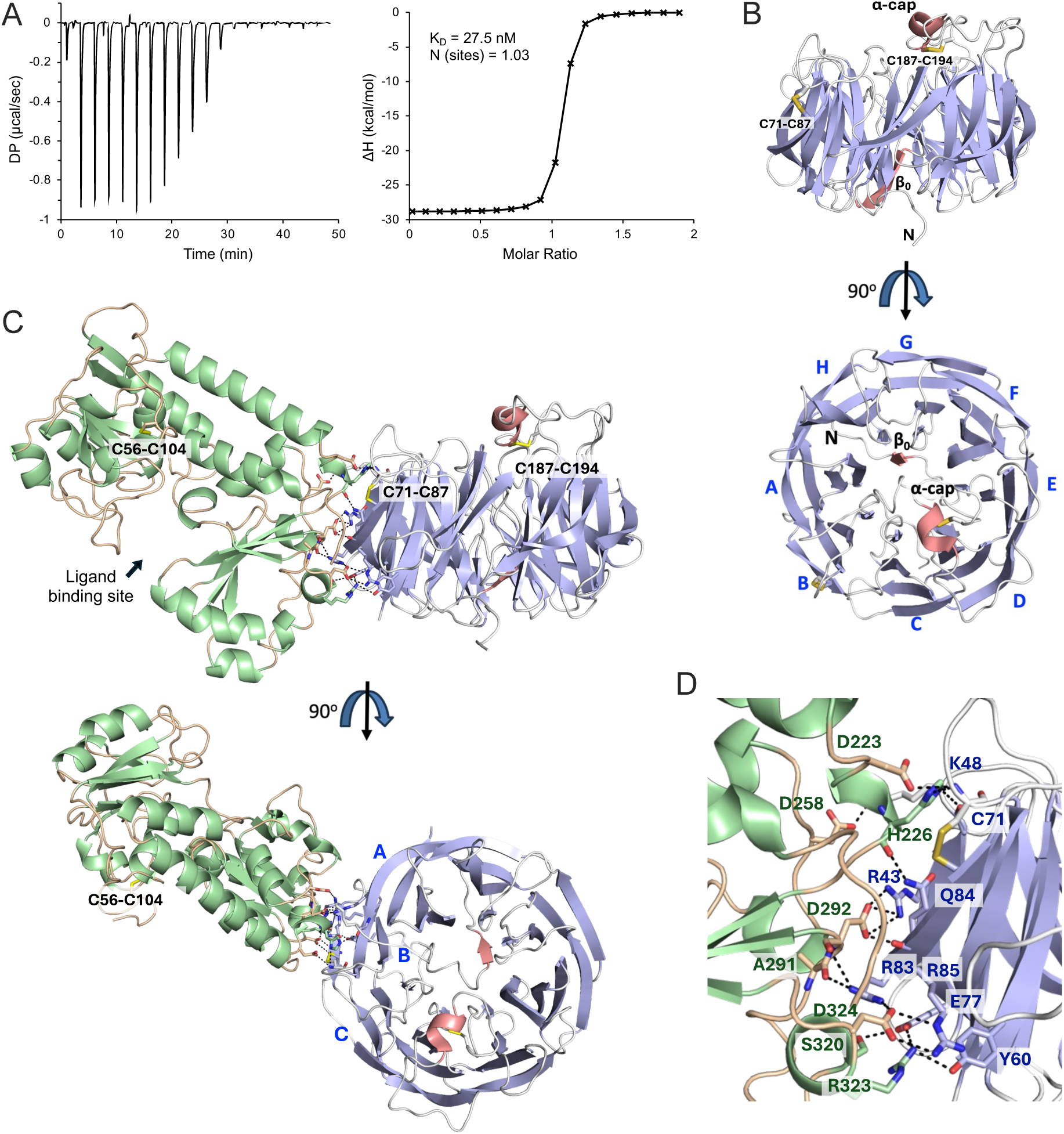
Binding affinity of FecB and Rv3035 and X-ray crystal structures of Mth Rv3035 and the FecB-Rv3035 complex. (A) A 19-point ITC experiment was carried out by titrating 200μM Mth Rv3035 into 20μM Mtb FecB. The resulting thermogram is shown (left graph), resulting in the following calculated parameters: K_D_ = 27.5 ± 4.6 nM; N(sites) = 1.03 ± 0.004 (right graph). (B) Cartoon depiction of the structure of Mth Rv3035. The β-propeller fold is colored in light purple for β-strands and white for loops, each blade was assigned a letter, and the additional secondary structure elements are colored in pink. The disulfide bonds are shown in stick with yellow sulfur atoms. (C) Cartoon depiction of the structure of the FecB-Rv3035 complex. Rv3035 is colored as in (B). FecB secondary structure elements are colored in light green and loops in wheat. (D) A zoom in of the interface of the FecB-Rv3035 complex with interacting residues shown in stick, and hydrogen bond and salt bridge interactions are represented in black dashed lines.

### Structure determination of Rv3035 and the FecB-Rv3035 complex

We next solved the structure of Mth Rv3035 by X-ray crystallography to 1.9 Å (table S1). Mth Rv3035 crystallized with two subunits in the asymmetric unit. In both subunits, there is clear electron density for all but the N-terminal seven residues of Rv3035. The two subunits are virtually identical with a root mean square deviation (rmsd) of 0.46 Å over 392 residues.

The structure of Rv3035 adopts a β-propeller fold, composed of eight 4-stranded β-sheets, where each β-sheet is referred to as a blade (Fig. 4B and fig. S4A). At the N-terminus of the protein, preceding the β-propeller fold, are ∼40 residues that form an ordered loop region, which occupies the bottom half of the central cavity formed by the β-propeller. A small portion of this N-terminal region forms a β-strand (β_0_, residues 27-32) that forms a β-sheet with the interior β-strand (β26) of blade H (Fig. 4B and fig. S4A). The top half of the cavity is occupied by a second ordered loop region formed by residues 171-197 containing a small α-helix (fig. S4A), which appears to act as a cap (α-cap) for the central pore (Fig. 4B).

Mth Rv3035 contains two disulfide bonds (Fig. 4B, fig. S4A, and fig. S6A). The first is formed between C71 and C87 and connects the exterior β-strand (β5) of blade B to the loop region on the α-cap side of the β-propeller, and the second is formed between C187 and C194 connecting the α-cap to the ordered loop region following it, perhaps stabilizing the cap region. A structural homology search of Mth Rv3035 utilizing the DALI server (*17*) (table S2) demonstrated that the closest hits have extremely high structural homology with Rv3035 and the majority are accessory proteins that assist in remodeling protein assemblies; for example, two of the top hits are BamB from Gram-negative bacteria.

To determine the protein-protein interacting residues of the FecB-Rv3035 complex, we solved the X-ray crystal structure of the FecB-Rv3035 complex to a resolution of 3.35 Å (table S1). There are minimal structural differences between FecB alone (*7*) and FecB in complex with Rv3035 (fig. S6, C and D). One notable difference is the presence of a disulfide bond between C56 and C104 (fig. S6D), which is absent in the previous FecB crystal structure. Likewise, there are little differences between the structures of Rv3035 alone and in complex with FecB (fig. S6, E and F).

The FecB-Rv3035 complex interface is small and involves 729 Å^2^ (5.6%) of the FecB surface, and 692 Å^2^ (4.3%) of Rv3035 (Fig. 4C). Although the interface is small, there are multiple interactions between the two proteins including 19 hydrogen bonds and salt bridges that are ≤ 3.5 Å and six interactions that are < 3.0 Å (Fig. 4D and table S3). The interface involves residues from Rv3035 blade B (β4-β5) and residues from the loop that precedes blade B (between β1 and β2) (Fig. 4D and fig. S4A). The interface also involves the backbone nitrogen atom from disulfide bond forming C71 (Fig. 4D and fig. S6B). Strikingly, only nine Rv3035 residues are responsible for the polar interactions at the interface, with six residues forming multiple contacts with FecB: R43, K46, E77, R83, Q84, and R85, where R83 forms four polar contacts at the interface. FecB residues participating in the polar contacts at the interface are localized to the second lobe of FecB and are located in α7 (D223, H226), the loop between α8 and α9 (D258), the loop between α10B and β11 (A291, D292), and α13 (S320, R323, D324)(*7*). Similar to Rv3035, several FecB residues are responsible for multiple contacts at the interface: H226, D292, S320, R323 and D324; where D292 forms five and D324 forms four polar contacts at the interface. The FecB-Rv3035 interface leaves the ligand binding site of FecB accessible to solvent, and thus still able to accommodate a ligand (Fig. 4C).

### Key FecB residues for binding Rv3035

To test the functional importance of FecB residues involved in binding Rv3035, we mutated FecB D292 to alanine alone or H226, D292 and D324 to alanine together, and tested binding of these proteins to Rv3035 by protein interaction pull-down assays and by ITC. After determining that the mutants were properly folded by circular dichroism (fig. S7A), we utilized FecB-His and untagged Rv3035 in a nickel-affinity pull-down assay (fig. S7B). While WT FecB-His copurified with Rv3035, FecB D292A alone and the FecB triple mutant H226A_D292A_D324A failed to pull down Rv3035. In addition, binding to Rv3035 was not observed for the single and triple FecB mutants by ITC analysis (fig. S7C). Thus, the nanomolar affinity (K_D_ of ∼30 nM) interaction between FecB and Rv3035 can be completely disrupted by mutating one FecB residue, D292.

### Rv3035 mediates intrinsic multidrug resistance in Mtb

Having demonstrated that FecB and Rv3035 interact both *in vivo* and *in vitro*, we next set out to probe the cellular function of Rv3035. In TnSeq screens, *rv3035* mutants are usually annotated as having a “growth defect” (*18*) and are often absent from TnSeq screen input libraries. That is likely the reason why *rv3035* was not a hit in our TnSeq chemical-genetic interaction screen that identified *fecB* as important for intrinsic antibiotic resistance in Mtb (*5*). Because *fecB* and *rv3035* display similar growth defects by CRISPRi (*19*), we reasoned that it would be possible to generate a *rv3035* knockout strain (Δ*rv3035*) to test directly whether Rv3035 mediates intrinsic resistance to different antibiotics. Although Δ*rv3035* colonies took longer to grow on agar plates than WT Mtb, the mutant strain exhibited similar growth kinetics as WT and Δ*fecB* in liquid media (fig. S8A). Additionally, we generated a Δ*fecB* Δ*rv3035* double knockout as well as the respective complemented mutants.

We performed MIC assays using these strains for several antibiotics having different mechanisms of action to which Δ*fecB* was hypersusceptible (Fig. 5A). Isoniazid was included as a control because its MIC is not affected by the absence of *fecB* (*5*). The single Δ*rv3035* and double Δ*fecB*Δ*rv3035* mutants showed very similar antibiotic hypersusceptibility patterns as Δ*fecB*, with reduced MICs compared to WT. As the basis for Δ*fecB*’s hypersusceptibility to antibiotics is likely a more permeable cell envelope compared to WT Mtb (*5*), we also tested whether Δ*rv3035* and Δ*fecB*Δ*rv3035* are more permeable to fluorescently labelled vancomycin and the reporter dye calcein-AM (*20*). Both mutants phenocopied Δ*fecB* with increased uptake of BODIPY-vancomycin (Fig. 5B) and both were hyperpermeable to calcein-AM similarly to Δ*fecB* (Fig. 5C).

**Figure 5.**
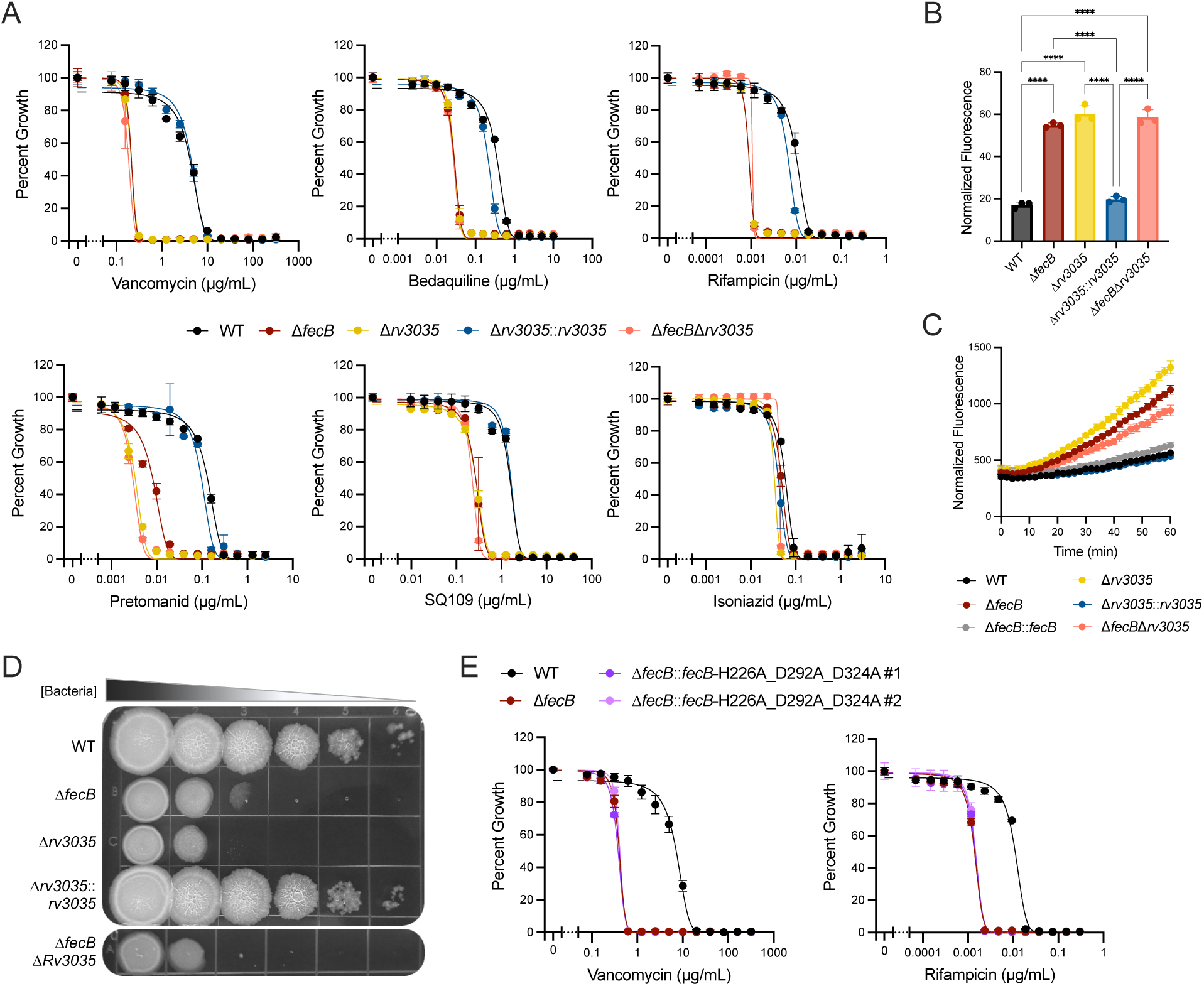
FecB and Rv3035 function in the same pathway that mediates cell envelope impermeability and antibiotic resistance in Mtb. (A) MIC assays for vancomycin, bedaquiline, rifampicin, pretomanid, SQ109 and isoniazid, percent growth calculated from no drug control wells (day 10). (B) BODIPY-vancomycin uptake after 30min of incubation, fluorescence intensity normalized by OD_580nm_. Statistical significance was determined by one-way ANOVA and Tukey post-hoc test. (C) Calcein-AM uptake, fluorescence intensity normalized by OD_580nm_ at 60min. (D) Spot Assay. (E) MIC assay for vancomycin and rifampicin, percent growth calculated from no drug control wells. Δ*fecB* complemented with FecB triple mutant that does not bind Rv3035 (Δ*fecB*::*fecB*-H226A_D292A_D324A) and control strains (day 14), #1-2 indicate individual colonies (biological replicates). Error bars: ±SD. \**P* < 0.05; \*\**P* < 0.01; \*\*\**P* < 0.001; \*\*\*\**P* <0.0001. N independent experiments = 2 (A-D), 1 (E), each performed with triplicate cultures.

Δ*fecB* also displayed altered colony morphology and a growth defect when growing as single colonies on agar plates, appearing smooth on high density spots and often failing to grow in lower spot dilutions (Fig. 5D). This makes experiments that rely on the quantification of Δ*fecB* colony forming unit (CFU) titers unreliable. Although we do not know the basis of this phenotype, it could be related to cell envelope alterations. Δ*rv3035* and Δ*fecB*Δ*rv3035* displayed the same defect (Fig. 5D). Taken together, these results demonstrate that Rv3035 is important for intrinsic multidrug resistance, mediates cell envelope impermeability and is required for regular growth on solid media. Importantly, deleting *fecB* and *rv3035* in the same strain did not result in enhanced phenotypes (Fig. 5, A-D), a result that it is expected for genes functioning in the same pathway.

To test whether binding between FecB and Rv3035 is required for maintaining cell envelope integrity, we complemented Δ*fecB* with an integrated plasmid expressing the FecB triple mutant H226A_D292A_D324A which fails to bind Rv3035 *in vitro* (fig. S7C). First, we confirmed that the FecB triple mutant is stably expressed *in vivo* (fig. S8B). Then, we tested whether Δ*fecB* complemented with the triple mutant (Δ*fecB*::*fecB*-H226A_D292A_D324A) can rescue Δ*fecB*’s hypersusceptibility to vancomycin and rifampicin. The MICs of these drugs were unchanged in Δ*fecB*::*fecB*-H226A_D292A_D324A compared to Δ*fecB* (Fig. 5E). Similarly, the FecB triple mutant failed to rescue Δ*fecB*’s growth defect in solid media (fig. S8C). These results indicate that FecB’s binding to Rv3035 is critical to mediate intrinsic drug resistance in Mtb.

### FecB and Rv3035 interact with cell envelope biosynthesis proteins

Our immunoprecipitation data revealed that FecB and Rv3035 also interact with the cell envelope biosynthesis proteins AftB and Rv0227c (Fig. 3, A and B), which was confirmed by co-immunoprecipitation assays (fig. S9 and S10). Moreover, *fecB* and *rv3035* are co-essential with *aftB* (Fig. 3C), which suggests they may perform similar functions and could be part of a protein complex (*12*). AftB catalyzes the last step in the biosynthesis of the arabinan domain of arabinogalactan (AG) and lipoarabinomannan (LAM), adding the terminal β-arabinofuranose (β-Ara*f*) residue to those molecules (*21*). In AG, the terminal β-Ara*f* and the penultimate α2-linked Ara*f* residue provide mycolic acid (MA) attachment sites which are critical for the formation of the mycolyl–arabinogalactan–peptidoglycan (mAGP) complex (*22*, *23*). Rv0227c (named polar growth factor A, or PgfA, in *Mycobacterium smegmatis*) is an inner membrane protein that interacts with MmpL3, the transporter for the cell envelope component trehalose monomycolate (TMM), and with a TMM analog, suggesting that it participates in TMM transport across the plasma membrane (*24*).

In the periplasm, TMMs may donate a mycoloyl residue to AG to form the mAGP complex or to another TMM molecule to form trehalose dimycolate (TDM or cord factor) (*25*). While MAs attached to AG (mAG) are the major components of the inner leaflet of the mycobacterial outer membrane (MOM), TMMs, TDMs and free MAs are thought to populate both the outer and inner leaflets of the MOM (*3*). These lipids are critical determinants of Mtb’s cell envelope impermeability (*3*). Since AftB and Rv0227c are involved in the biosynthesis of mAG and the MOM, the interaction of FecB and Rv3035 with these proteins suggests that they participate in the assembly or maintenance of mAG and/or the MOM.

We used AlphaFold 3 (*26*) to predict the potential protein complexes formed by FecB, Rv3035, AftB and/or Rv0227c, modeling different combinations of these proteins (fig. S11 and S12). The FecB and Rv3035 complex showed a very high per-residue model confidence score (plDDT, fig. S11), and has the same interface and orientation observed in the experimental structure of FecB-Rv3035, superimposing with a rmsd of 0.91 Å over 696 residues (Fig. 4C). The complexes containing AftB (with or without Rv0227c) showed a higher confidence score when modeled with Rv3035 than with FecB alone (fig. S12, B and C). These structural models crossed the interface predicted template modeling (ipTM) score threshold for high-quality prediction of 0.8 when AftB was modeled with FecB and Rv3035, with Rv3035 and Rv0227c, or when all 4 proteins were modeled together (fig. S12). This suggests that Rv3035 bridges the interaction between FecB and AftB, and between FecB and Rv0227c.

### *AftB* is more vulnerable to knockdown in Δ*fecB* than in WT

Because *aftB* shares an essentiality profile with *fecB* and *rv3035* (Fig. 3C), we decided to further explore the genetic interaction between *fecB* and *aftB.* We depleted *aftB* with a weak CRISPRi single guide RNA in both WT and Δ*fecB*. While this weak guide caused a slight growth defect in WT upon induction of the CRISPRi system, knockdown of *aftB* by the same weak guide in Δ*fecB* completely abrogated growth (Fig. 6A). This indicates that *aftB* is more vulnerable to knockdown in Δ*fecB* than in WT as a moderate depletion of *aftB* leads to a profound loss of fitness in the absence of *fecB*.

**Figure 6.**
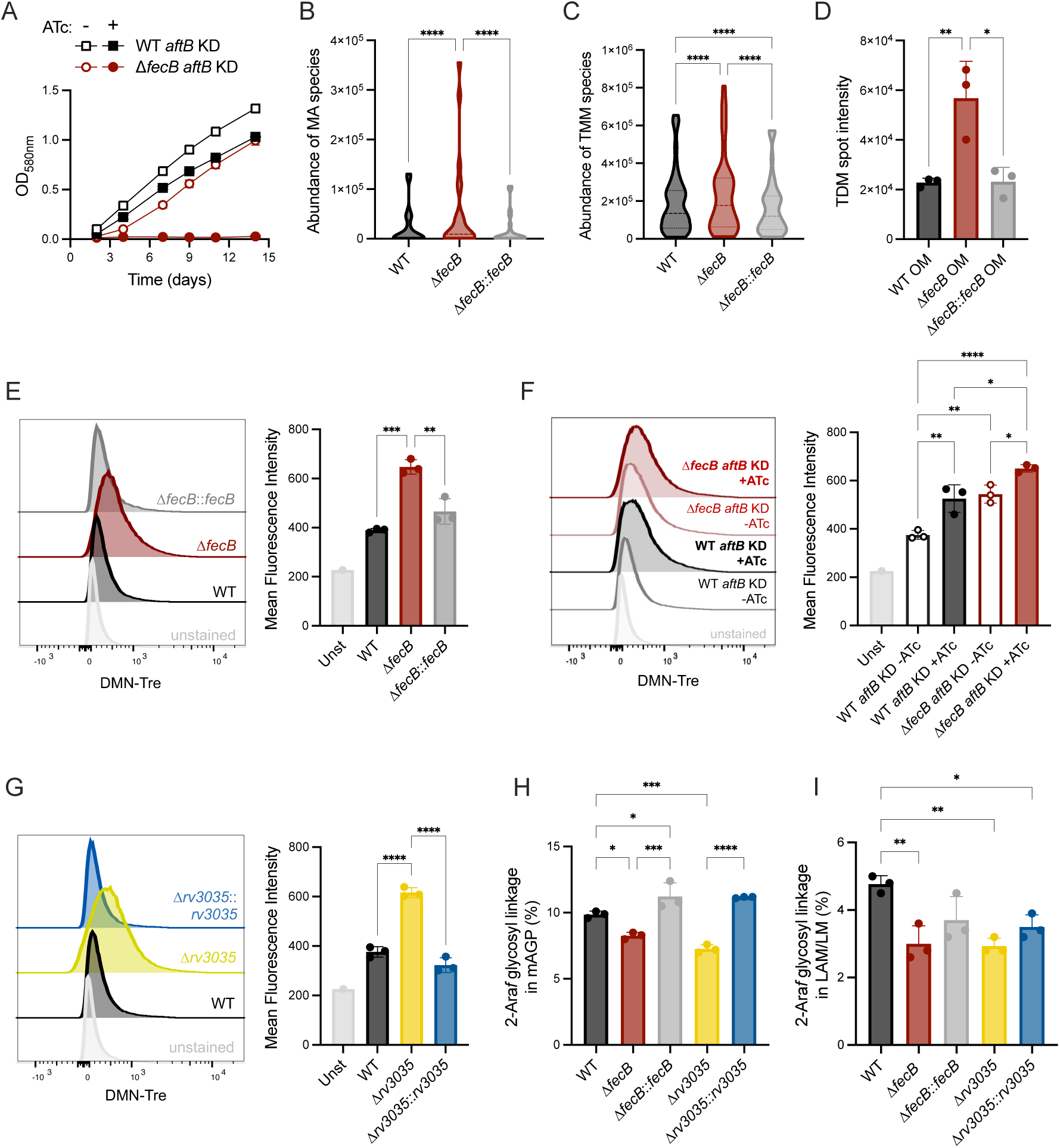
FecB and Rv3035 support AftB’s function. (A) Growth curves of CRISPRi KD of *aftB* in WT/Δ*fecB* in the absence or presence of ATc inducer; strains were grown in 384-well plate in standing conditions. (B-C) Lipidomic analysis by LC-MS of total lipids identifying (B) free-mycolic acid (MA) species and (C) trehalose monomycolate (TMM) species. (D) Trehalose dimycolate (TDM) spot intensity quantified from fig. S13I by densitometry analysis using ImageJ software. OM refers to the outer leaflet of the outer membrane obtained after AOT treatment. (E-G) Flow cytometry analysis of DMN-Tre labelled cells using 405nm laser and 525/50 channel. Representative histograms (left) and quantification (right) of mean fluorescence intensity of DMN-Tre probe. Unstained (Unst) WT was used as a control for autofluorescence. (H-I) Relative percentage of 2-arabinofuranosyl (2-Ara*f*)-linked residues in (H) mAGP and (I) lipoglycans (LAM and LM mixture). See fig. S14 for full details of the glycosyl linkage analysis of mAGP and LAM/LM. Statistical significance was determined by two-way (B-C) or one-way (D-I) ANOVA and Tukey post-hoc test. Error bars: ±SD. \**P* < 0.05; \*\**P* < 0.01; \*\*\**P* < 0.001; \*\*\*\**P* <0.0001. N independent experiments = 2 (A, E-G), 1 (B-D, H-I), each performed with triplicate cultures.

### FecB, Rv3035 and AftB are required to maintain normal TMM levels in Mtb

To test whether FecB is required for proper assembly of the MOM, we performed lipidomics analysis of total extractable lipids from WT, Δ*fecB* and Δ*fecB*::*fecB* strains (Fig. 6, B and C, and data S4). Contrary to expectations that mycolate levels would be reduced in Δ*fecB* due to their role in mediating envelope impermeability, the mutant displayed a higher abundance of TMMs and free MAs compared to WT and Δ*fecB*::*fecB*.

Since TMMs accumulate in the plasma membrane of TMM transport mutants (*24*, *27–29*), we next assessed if lipid trafficking was disrupted in Δ*fecB*. We treated bacterial pellets with the anionic surfactant dioctylsulfosuccinate sodium (AOT) dissolved in heptane to strip non-covalently bound lipids off the MOM (*30*). In *Corynebacterium glutamicum*, AOT in heptane forms reverse micelles which solubilize and capture OM surface lipids but does not penetrate other layers of the cell envelope (*30*). We validated this approach by analyzing known lipids extracted from the supernatants and pellets of AOT treated samples, corresponding to the outer leaflet of the MOM (labelled OM) and all other extractable lipids (labeled inner membrane or IM), respectively (fig. S13, A-D, and data S4). Δ*fecB* displayed a higher abundance of TMMs and free MAs in both IM and OM fractions, compared to WT and Δ*fecB*::*fecB* (fig. S13, E-H, and data S4). We used the OM fraction to visualize and quantify TDMs via thin-layer chromatography (Fig. 6D and fig. S13I). Although TDM levels are inversely correlated with TMMs in mutants with defective TMM transport (*24*, *27–29*), Δ*fecB* also displayed increased TDM levels compared to controls.

To validate the lipidomics data with an alternative method, we used the solvatochromic trehalose probe DMN-Tre to track MA incorporation into TMMs via flow cytometry (*31*). Upon entering the cell, DMN-Tre is mycolylated to form DMN-Tre monomycolates, which are subsequently inserted into the MOM, where the hydrophobic environment activates the probe’s fluorescence (*31*). Corroborating the lipid analysis, Δ*fecB* incorporated more DMN-Tre in its cell envelope than WT, shown as increased mean fluorescence intensity compared to WT and Δ*fecB*::*fecB* (Fig. 6E). The same phenotype was observed in *aftB* KD strains upon CRISPRi activation (Fig. 6F) and in Δ*rv3035* compared to controls (Fig. 6G). These results indicate that *fecB*, *rv3035* and *aftB* mutants accumulate trehalose monomycolates in the MOM.

Overall, these data suggest that FecB, Rv3035 and AftB are important for proper assembly and/or maintenance of free mycolates in the MOM.

### FecB and Rv3035 are important for arabinofuranosyltransferase function in Mtb

To investigate whether the FecB-Rv3035 complex influences AftB’s arabinofuranosyltransferase activity, we quantified per-*O*-methylated alditol acetates derived from the mAGP complex and lipoglycans (LAM and LM mixture) from WT, Δ*fecB*, Δ*rv3035* and complemented strains. 2-linked arabinofuranosyl (Ara*f*) residues in the mAGP and LAM were reduced in Δ*fecB* and Δ*rv3035*, compared to WT (Fig. 6, H and I). This defect was fully reversed in the mAGP and partially reversed in the LAM of the complemented strains (Fig. 6, H and I). These findings are consistent with reduced AftB activity in Δ*fecB* and Δ*rv3035* (*21*, *32*). The greater decrease in 2-Ara*f* linkage levels in LAM (1.6-fold) compared to AG (1.2 and 1.3-fold) of the two mutants, and the more efficient restoration of these glycosyl linkages in AG upon genetic complementation, indicate that the available AtfB activity in each strain prioritizes the completion of the arabinan domain of AG (which is critical to the formation of the MOM) over that of LAM.

Noticeably, in mAGP (fig. S14, A-H) the decrease in 2-Ara*f* linkages in the mAGP of Δ*fecB* and Δ*rv3035* was accompanied by a modest decrease in 5-Ara*f* linkages and a reduction in overall Ara*f*/Gal*f* ratio in Δ*rv3035,* suggestive of a smaller size arabinan domain compared to WT and Δ*rv3035*::*rv3035* strains (fig. S14, B and H). With regards to LAM (fig. S14, I-O), the decrease in 2-Ara*f* linkages in Δ*fecB* and Δ*rv3035* was accompanied by a decrease in branched 3,5-Ara*f* linkages and *t*-Ara*f* (fig. S14, I and K) and minor changes in the mannan domain of the lipoglycans (fig. S14, L-O), except for a marked increase in 6-mannopyranosyl (6-Man*p*) residues in Δ*fecB* compared to WT, and in 2,6-Man*p* and 2-Man*p* residues in Δ*rv3035* compared to WT and Δ*rv3035*::*rv3035*, typifying a more branched mannan domain in LM and/or LAM (fig. S14, M-O). Collectively, these structural alterations in the AG and lipoglycans of Δ*fecB* and Δ*rv3035* point to greater alterations in the biosynthetic machinery of these two glycans than a sole reduction in AftB activity and suggest that other arabinosyltransferases and mannosyltransferases may be negatively impacted by the loss of FecB or Rv3035.

### FecB is essential for virulence of Mtb in mice

The mycobacterial cell envelope is a formidable barrier that protects Mtb not only from antibiotics but also from host immune responses (*3*). Thus, we hypothesized that *fecB* might also be required for Mtb to establish a successful infection in mice. The unreliability of Δ*fecB* to grow as single colonies on agar plates (Fig. 5D), however, impairs the quantification of the bacterial burden in infected mice. To circumvent this problem, we implemented a genetic switch to activate *fecB*’s expression in the presence of a tetracycline inducer (TetON) (*33*) and allow quantification of CFUs from mice infected with Δ*fecB* harboring the TetON switch (FecB-TetON). The conditional expression of *fecB* was validated via MIC assays with vancomycin and rifampicin, which showed that addition of ATc rescued the antibiotic hypersusceptibility of FecB-TetON (Fig. 7A).

**Figure 7.**
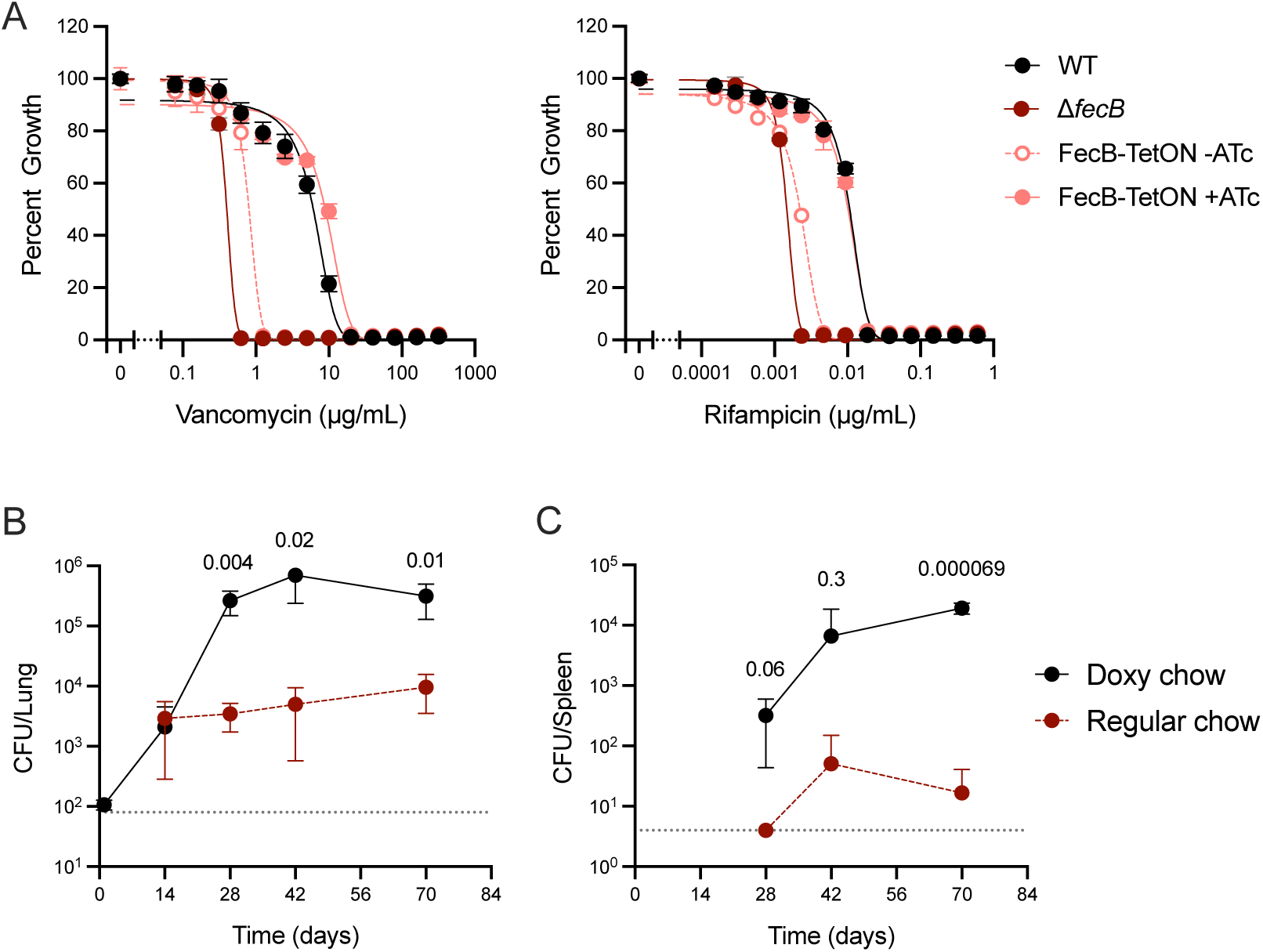
FecB is required for Mtb virulence. (A) MIC assays for vancomycin and rifampicin, in the absence or presence of ATc to induce *fecB* expression in the FecB-TetON strain (day 14). Percent growth calculated from no drug control wells. (B-C) Growth and persistence of FecB-TetON in the (B) lungs and (C) spleens of mice receiving doxy or regular chow. Data are average CFU counts from four mice per time point. Dashed black lines represent lower limit of detection. Statistical significance was determined by unpaired T-tests for each time point, p-value is shown above each pair analyzed. Error bars: ±SD. N independent experiments = 1 performed with triplicate cultures (A), 1 (B-C).

Mice were infected via the aerosol route with FecB-TetON and received either regular or doxycycline (doxy) chow throughout the course of the experiment (Fig. 7, B and C), with doxy chow inducing *fecB* expression. Lung and spleen homogenates from both groups were cultured on agar containing ATc to activate *fecB* expression *ex vivo*. FecB-TetON grew and established a high titer chronic infection in the lungs of mice fed with doxy-containing chow (Fig. 7B). In contrast, in mice receiving regular chow, in which expression of *fecB* is silenced, FecB-TetON grew during the first 14 days post-infection (p.i.), after which CFU counts remained relatively stable for another 8 weeks p.i. Dissemination to the spleen was delayed, and similarly to the lung, spleen bacterial burden was significantly lower in mice receiving regular versus doxy chow (Fig. 7C). Thus, we conclude that *fecB* is required for Mtb virulence and plays a critical role during growth in the acute phase of infection but is not required for Mtb’s persistence in the host.

## Discussion

The mycobacterial cell wall, comprised of peptidoglycan, arabinogalactan and mycolic acids, has been a key target of several anti-TB drugs (*34*). Two of the four first-line TB drugs, isoniazid and ethambutol, target mycolic acid and arabinogalactan biosynthesis, for instance. Although these drugs were developed more than 60 years ago, they remain part of the standard TB chemotherapy due to their success in treating drug-susceptible Mtb infections, which validates efforts to explore novel inhibitors that target the cell wall of Mtb (*10*). Despite being an important target for drug development and playing a central role in Mtb pathogenesis and intrinsic drug resistance, many aspects of the Mtb cell wall, and more broadly, cell envelope, such as factors involved in their assembly and maintenance remain unknown. In the current study, we describe two periplasmic proteins, FecB and Rv3035, that work together to support arabinofuranosyltransferase activity, mediate cell envelope impermeability and impart intrinsic multidrug resistance in Mtb.

Our transcriptomic data indicate that Mtb lacking *fecB* upregulates the *iniBAC* operon (**i**so**n**iazid **i**nduced genes B, A and C), which is named for being induced by isoniazid, ethambutol, and other inhibitors of cell wall biosynthesis (*10*). Upregulation of the *iniBAC* genes in Δ*fecB* occurred independently of the presence of detergent in the media and most likely reflects a constitutive defect in cell wall integrity, which results in Δ*fecB’s* increased membrane permeability (*5*).

The iron deprivation transcriptional signature and accumulation of siderophores in Δ*fecB* grown in detergent-free media points to *fecB* having an activity in iron homeostasis. However, whether this is a direct or indirect effect remains unclear. While FecB can bind ferric-carboxymycobactin with high affinity *in vitro* (*7*), experiments probing the requirement of *fecB* for iron acquisition *in vivo* showed only minor growth phenotypes. FecB variants with lower binding affinity to ferric-carboxymycobactin (FecB E339S_R240S and E339S_Y242S) rescued the iron deprivation transcriptional signature and slower growth of Δ*fecB* in iron-limiting conditions, suggesting that binding to iron-loaded siderophores is dispensable in FecB’s ability to reverse these phenotypes. On the other hand, FecB E339S_R240S and E339S_Y242S have slightly higher binding affinity to apo-carboxymycobactin and thus we could not test the *in vivo* requirement of FecB’s binding to iron-free apo-siderophores using these variants. Importantly, using FecB E339S_R240S and E339S_Y242S, and other genetic and microbiological approaches, we showed that FecB’s role in iron homeostasis is likely not connected to its role in mediating antibiotic resistance in Mtb.

Specific evidence for FecB’s function in cell envelope integrity came to light upon identification of FecB’s most enriched physical interactors: Rv3035, AftB and Rv0227c. In *Mycobacterium marinum*, FecB was also found to interact with AftB and the Rv3035 homologue MMAR_1667 (*35*). Although AftB’s function has been well described in the literature (*21*, *32*, *36*), only a few studies explore the function of Rv0227c (*24*, *37*) and Rv3035 (*35*) homologues in mycobacteria. Here we report on the structure of Rv3035 alone and of the FecB-Rv3035 complex, which identified key FecB residues involved in binding Rv3035 that we were able to validate experimentally. Moreover, we show that FecB’s binding to Rv3035 is required for its function in mediating antibiotic resistance in Mtb. MtbΔ*rv3035* and the double knockout Δ*fecB*Δ*rv3035* displayed the same phenotypes indicative of perturbed cell envelope integrity as Δ*fecB*. This indicates that the absence of either protein equally impacts the function of the complex and thus both are required to fulfill their function.

Structural homology analysis for Rv3035 revealed two classifications of structural homologs (table S2). One class represents assembly factors that contribute to scaffolding in larger complexes, facilitating the coordinated movement of multiple protein components. This includes BamB of the prokaryotic β-barrel assembly machinery (BAM) complex, and the eukaryotic ODA16, an assembly factor involved in outer dynein arm intraflagellar transport (*38*). When comparing these structural homologs to Rv3035, both the N-terminal propeller cavity plug region and the α-cap present in Rv3035 are absent in these assembly factors. The second class of Rv3035 structural homologs have enzymatic activity. Strikingly, these structural homologs - *Acidithiobacillus ferrooxidans* tetrathionate hydrolase (*39*), *Methylacidiphilum fumariolicum* SolV methanol dehydrogenase (*40*), and the *Devosia albogilva* quinone-dependent dehydrogenase (*41*) - all have secondary structure elements similar to the Rv3035 α-cap and bear an N-terminal propeller central cavity plug. However, in these enzymatic structural homologs, the propeller fold is more extensively decorated than in Rv3035. Indeed, Rv3035—due to the presence of the N-terminal propeller cavity plug and the propeller decorations—has greater similarity with its enzymatic structural homologs than it does with the assembly factor homologs. Although our studies show that Rv3035 acts as a scaffolding protein in its interactions with FecB, its resemblance to the enzymatic homologs raises the possibility that Rv3035 may possess an as-yet unidentified enzymatic activity.

What is the function of FecB and Rv3035, and how do these proteins contribute to intrinsic multidrug resistance in Mtb? Our findings suggest that FecB and Rv3035 interact with AftB, a known arabinofuranosyltransferase, and together form a functional complex that supports cell envelope biosynthesis. Proteomic data, CRISPRi essentiality profiles, and high-confidence structural predictions from AlphaFold 3 collectively support the interaction of FecB-Rv3035 with AftB. In the predicted model, Rv3035 acts as a bridge between FecB and AftB, positioning the periplasmic proteins in close proximity to AftB’s active site cavity (*42*). The proposed orientation suggests that the FecB-Rv3035 complex may serve as scaffold for AftB’s engagement with its substrate AG/LAM or may regulate AftB’s enzymatic activity.

Lipidomics data further support FecB’s role in AftB function as Δ*fecB* exhibited increased levels of TMMs, TDMs and free MAs compared to controls. This phenotype is observed in *M. smegmatis* treated with ethambutol (*43*) (inhibitor of AG/LAM biosynthesis), and in deletion mutants of *aftB* (*21*) and *aftC* (*44*) (another arabinofuranosyltransferase) in *C. glutamicum* and *M. smegmatis*, respectively, all of which show increased levels of trehalose mycolates. Notably, *fecB*, *rv3035* and *aftB* mutants all displayed enhanced incorporation of TMMs into the MOM, a phenotype exacerbated by selective depletion of *aftB* in the Δ*fecB* background. This depletion completely inhibited growth in Δ*fecB*, while only mildly affecting WT, indicating that Mtb becomes more reliant on full *aftB* expression in the absence of FecB.

Further evidence of FecB and Rv3035’s functional support of AftB comes from glycosyl linkage analysis. Mtb mutants lacking *fecB* or *rv3035* showed reduced levels of 2-linked Ara*f* residues in AG and LAM, consistent with impaired terminal Ara*f* addition by AftB (*21*, *32*). A decreased ratio of arabinose to galactose in AG was also observed in a *M. marinum fecB* mutant (*35*), supporting our findings of additional alterations in length and branching of AG and LAM. Besides the impact on mycolate levels and their localization, structural alterations in AG and LAM may compromise overall cell envelope integrity and influence host-pathogen interactions (*32*, *35*, *45*).

Taken together, our data support a model in which the FecB-Rv3035 complex is essential for optimal AftB activity *in vivo*. This complex contributes to the biosynthesis and structural integrity of the mycobacterial cell envelope, thereby playing a key role in Mtb’s intrinsic resistance to antibiotics. This new understanding of the molecular mechanisms that govern Mtb’s cell envelope impermeability and intrinsic antibiotic resistance provides valuable insights for TB drug discovery and the development of more effective regimens that may shorten TB treatment.

## Materials and Methods

### Bacterial strains and culture conditions

Mtb strains used in this study are listed in data S5. For most experiments, the strains were cultured in Middlebrook 7H9 medium (BD, #271310) supplemented with 0.2% glycerol, 0.5% bovine serum albumin fraction V, 0.2% dextrose, 0.085% NaCl and 0.05% Tween80 (Sigma, # P8074), this is referred to as “regular” medium. For detergent-free conditions, Tween80 was omitted from the media recipe. For growth in iron-limiting conditions, self-made 7H9 medium without ammonium ferric citrate (AFC) was prepared and chelated for 2 days with 20g/liter of BioRad Chelex 100 resin. After filter-sterilization, metals other than iron were added according to their concentrations in Middlebrook 7H9. To prepare Low/High Iron media, 1µM or 160µM of AFC was added to iron-chelated medium, respectively. For reference, Middlebrook 7H9 medium contains 150µM AFC. Where required, antibiotics or small molecules were used at the following concentrations: hygromycin (50μg·ml^−1^), kanamycin (25μg·ml^−1^), zeocin (25μg·ml^−1^), streptomycin (25μg·ml^−1^), apramycin (60μg·ml^−1^) and ATc (200ng·ml^−1^ for CRISPRi strains and 1000ng·ml^−1^ for FecB-TetON). The strains were grown at 37°C and 5% CO_2_ in a standing incubator, unless otherwise indicated.

### Plasmids construction

Plasmids used in Mtb strains are listed under the strain “full name” after the double colon in data S5. Plasmids starting with pGM were constructed using Invitrogen Gateway Cloning system. Constructs harboring the gene of interest were synthesized by GenScript. CRISPRi constructs were generated as previously described (*19*).

### RNA extraction

Total RNA was extracted from 10mL of mid-log phase bacterial cultures by first adding 1:1 guanidinium thiocyanate buffer followed by centrifugation for 10 minutes at 4°C. Bacterial pellets were bead-beaten into 0.9mL Trizol, 0.2mL chloroform was added to each sample and RNA in the aqueous phase was purified using the RNA Clean and Concentrator-25 kit (Zymo Research #R1017). After elution, samples were incubated with TURBO DNase (Ambion, AM2238) at 37°C for 1h to remove residual genomic DNA. Samples were then repurified using the RNA Clean and Concentrator-25 kit or RNA Clean and Concentrator-5 kit (Zymo Research #R1015).

### RNA-sequencing (RNA-seq) and analysis

Ribosomal RNA (rRNA) was removed from total RNA as previously described (*46*). The resulting rRNA-depleted RNA was then concentrated with the RNA Clean and Concentrator-5 kit (Zymo Research #R1015). RNA-seq libraries were created from these samples using the NEBNext Ultra II Directional RNA Library Prep Kit for Illumina (New England Biolabs, E7760L) along with NEBNext Multiplex Oligos for indexing (New England Biolabs). The libraries were purified using Dynabeads MyOne Streptavidin C1 (Thermo Fisher #65002), assessed for quality with a Bioanalyzer, and pooled based on their unique indexes. Sequencing was carried out on an Illumina NextSeq2000 platform at the Genomics Resources Core Facility of Weill Cornell Medicine.

RNA-seq data analysis was conducted using R. Sequencing reads underwent trimming and filtering with Rfastp, followed by alignment to the H37Rv reference genome (GenBank AL123456.3; RefSeq NC_000962.3; GCF_000195955.2; Genome assembly ASM19595v2) using the Rsubread package (*47*, *48*). BAM alignment files were generated, sorted, and indexed using Rsamtools. Read quantification was carried out with featureCounts, and differential gene expression analysis was performed using DESeq2 (*49*, *50*).

### Lipid extraction

Mtb cultures were grown in detergent-free 7H9 containing fatty acid-free BSA until mid-log phase. Bacterial pellets were transferred to sterile filters (*51*) which were added directly to 16 x 100mm glass tubes containing 2:1 chloroform: methanol or to 2mL screw cap tubes containing 1mL of 10mM AOT dissolved in heptane or PBS as a control. For AOT treatment, pellets in AOT or PBS were agitated in a bead-beater at low speed (2800rpm) for 10s to knock the cells off the filters followed by removal of filters and incubation in a cooling rack for 1h, with mixing by inversion of the tubes every 5 minutes. 500µL of AOT supernatant was added to 16 x 100mm glass tubes containing 2:1 chloroform: methanol. AOT and PBS pellets were resuspended in 600µL of 2:1 chloroform: methanol and added to 16 x 100mm glass tubes containing 2:1 chloroform: methanol.

### Liquid chromatography-mass spectrometry (LC-MS) analysis

Lipid extracts were centrifuged at 4000rpm for 15 minutes at 4°C and the supernatant decanted into new 16 x 100mm borosilicate tubes. The tubes were dried using a GeneVac evaporator under vacuum with gentle heating to 40°C. The residual residue was transferred to pre-weighed 4mL amber vials using 2mL 1:1 chloroform: methanol. The solution was again dried using the GeneVac evaporator. The vials were re-weighed to obtain the dry weight of lipid extract, and this mass was used to prepare a 1mg/mL solution in 1:1 chloroform: methanol. Lipids were stored at -20°C as a stock solution. A 150µL aliquot of stock solution was transferred to a glass insert tube (250µL, Agilent Technologies) nested within a microfuge tube. The aliquot was evaporated to dryness using an argon air stream then resuspended in 150µL of a solution of 70% hexanes and 30% isopropyl alcohol. After briefly vortexing to resuspend, the samples were centrifuged at 4000rpm for 15 minutes at 4°C. A volume of 100µL of the supernatant was transferred to a 2mL amber vial (Agilent) equipped with 250µL glass insert tubes and capped with 9mm septum-embedded screw caps (Agilent). A 10µL injection of this sample was analyzed using liquid chromatography mass spectrometry (LC-MS) on an Agilent 6520 ESI-QTOF. Separation of lipids using a 3µm diol column (2.1 x 150mm) and gradient elution from solvent A (70% hexane, 30% isopropanol, .02% formic acid, .01% ammonium hydroxide) to solvent B (70% isopropanol, 30% methanol, .02% formic acid, .01% ammonium hydroxide) as previously described (*52*). Following chromatography, the sample underwent electrospray ionization at 325°C with 5 L/min drying gas flow at 5500 V and 30 psig nebulizer pressure and ions were detected by time-of-flight in both positive and negative modes (m/z range: 100 – 3000). Internal calibrants (922.009798 positive mode, 983.0325 negative mode) were infused continuously.

Lipids were identified using unique mass-retention time identifiers for masses exhibiting the expected isotopic distributions as previously reported (*52*). Integrations of ion abundance were determined using Agilent Profinder 8.0 and Qualitative Analysis 6.0.

### Growth curves

Mtb strains were inoculated from frozen glycerol stocks and grown with appropriate selection antibiotics in 7H9 medium until mid-log phase at 37°C.

For pre-depletion of iron internal stores, bacteria were washed in PBS or iron-chelated 7H9 and diluted to an OD_580nm_ of 0.05 in iron-chelated 7H9. After 4-5 generations, bacteria were spun down at 800rpm for 8 minutes to generate a single cell suspension. Bacteria were diluted to an OD_580nm_ of 0.025 in the indicated media and added to flasks which were incubated standing at 37°C. For serial passaging, when cultures in high iron media reached stationary phase, low iron cultures were spun down at 800rpm for 8 minutes to generate a single cell suspension and bacteria were diluted to an OD_580nm_ of 0.025 in both high/low iron media. OD measurements from cultures in flasks were performed in a spectrophotometer. For growth curves in low iron media without detergent, iron pre-depleted cultures were washed in PBS or detergent-free iron-chelated 7H9, diluted to an OD_580nm_ of 0.05 and 100µL of bacteria were added to a 96-well plate in triplicates. OD measurements from plates were performed in a plate reader. Cultures grown in 96-well plates were resuspended prior to OD_580nm_ readings.

For CRISPRi strains, cultures were split into two flasks and 200ng/mL of ATc added to one of the flasks 2-3 days prior to setting up the growth curve to pre-deplete the gene of interest. Fresh ATc (200ng/mL) was again added to the media (+ATc conditions) before distributing bacteria into 96/384-well plates in triplicates.

### Reverse transcription quantitative polymerase chain reaction (RT qPCR)

Primers and probes for RT qPCR are listed in table S4. Primers and probes were ordered from TIB MOLBIO, LLC. cDNA was generated using MluV reverse transcriptase (New England Biolabs) and approximately 500ng of total RNA. Parallel no-reverse transcriptase reactions were prepared as a control for residual genomic DNA. qPCR was performed in the LightCycler® 480 System (Roche). The housekeeping gene *sigA* (Rv2703) was used as an internal control. *sigA* probes were labeled with 5’ LC760/3’ BBQ. Probes targeting genes of interest were all labeled with 5’ 6FAM/3’ BBQ or 3’ BHQ.

### Minimal inhibitory concentrations (MIC) assays

Mtb strains were inoculated from frozen glycerol stocks and grown with appropriate selection antibiotics in 7H9 medium until mid-log phase at 37°C. Bacteria were diluted to an OD_580nm_ of 0.01 in 7H9 and 50µL of bacteria were added to a 384-well plate in triplicates per antibiotic concentration, or to triplicate control wells without antibiotics. The HP D300e digital dispenser was used to deliver antibiotics to plate wells. Bacteria were grown in standing cultures at 37°C and OD_580nm_ was measured in a plate reader between days 10-21 following treatment (time point is indicated in each figure legend). For MICs in low iron or detergent-free conditions, bacterial cultures were first washed in PBS once or twice prior to dilution into the respective media of the MIC assay. For CRISPRi strains, cultures were split into two flasks and 200ng/mL of ATc added to one of the flasks 2-3 days prior to setting up the MIC assay to pre-deplete the gene of interest. Fresh ATc (200ng/mL) was again added to the media of the assay (+ATc conditions) before distributing bacteria into plates.

### Immunoprecipitation tandem mass spectrometry

#### Sample preparation

Whole-cell lysates were collected from log-phase cultures grown in the shaking incubator, incubated with 1% n-dodecyl -D-maltoside for 2h on a rotator at 4°C, followed by incubation with anti-DYKDDDDK (FLAG) magnetic agarose (Pierce #A36797). Captured proteins were dissociated from magnetic agarose by addition of 3x DYKDDDDK (3xFLAG) peptide (Pierce #A36805), according to the manufacturer’s instructions. Sodium Dodecyl Sulfate (SDS) was added to the peptide eluate at a final concentration of 2% and samples were boiled at 95°C for 10 minutes. Boiled samples were transferred to protein lo-bind tubes. FLAG- (or 3xFLAG-) tagged bait proteins were detected in eluates on an SDS-Polyacrylamide Gel Electrophoresis (PAGE) gel prior to mass spectrometry.

23µg of proteins per sample were flash frozen on dry ice and subsequently dried using a Speed-Vac. Next, 23µl of lysis buffer (5% SDS, 50mM TEAB) was added to initiate the S-Trap™ Micro Spin digestion protocol (PROTIFI). The proteins underwent reduction by 4mM dithiothreitol (DTT) and alkylation by 20mM iodoacetamide. Following an overnight digestion with trypsin at 37 °C, peptides were eluted first by addition of 40µL of 50mM triethylammonium bicarbonate (TEAB) and spun at 4000 rpm for 1 min. This was followed by a 1-minute spin with 40µL of 0.2% formic acid (FA) and, lastly, 40µL of 50% acetonitrile (ACN). The three elution were then pooled together, dried using a Speed-Vac, and reconstituted in 20µL of 0.1% FA in 5% ACN. After reconstitution, the samples were centrifuged at 16,000 rpm for 16 min and an 18µL aliquot of the supernatant was transferred to a non-binding HPLC vials.

#### LC-MS/MS and data analysis workflows

Data from the first experiment (pull-down of FecB-FLAG) was acquired on a TimsTOF Pro2 (Bruker) mass spectrometer, which was coupled to a nanoElute LC system from Bruker. 5µL of peptides were loaded and separated on an in-house 75 μm I.D. fused silica analytical column packed with 25-cm ReproSil-Pur C18-AQ (Dr. Maisch, GmbH, 120 Å, 3 μm) particles to a gravity-pulled tip, using an aqueous mobile phase (A) of water and 0.1% FA and an organic mobile phase (B) of ACN and 0.1% FA. A 30-minute gradient was employed for DDA-PASEF with a flow rate set at 500 nL/min. The captive nano-electrospray voltage was maintained at 1600V, using one column configuration (no trap). The DDA-PASEF method consists of 10 MS/MS PASEF scans per topN acquisition cycle, with ramp and accumulation times of 100 ms, covering an m/z range from 100 to 1700 and an ion mobility range (1/K0) from 0.70 to 1.30 V s/cm^2. Collision energy settings followed a linear function of ion mobility, ranging from 20 eV at 0.6 V s/cm^2 to 59 eV at 1.6 V s/cm^2, using default parameters. Calibration of the instrument was performed using three ions from the ESI-L Tuning Mix (Agilent) (m/z 622, 922, 1222).

The custom workflow ‘LFQ-MBR’ from FragPipe version 21.1 was used, including database search with MSFragger (version 4.0), deep-learning prediction rescoring with MSBooster, Percolator and ProteinProphet (Philosopher version 5.1.0) for PSM validation and protein inference. The raw files from TimsTOF Pro2 were searched against the database “Mycobacterium_tuberculosis_H37Rv_v4modified” appended with common contaminant proteins. Decoy reversed sequences were appended to the search database. The default MSFragger search parameters were used, except precursor and fragment mass tolerances were set to 20 ppm; enzyme: trypsin (not cutting before P); peptide length 7-30. Variable modifications were set to oxidation of Met and acetylation of protein N-t, and fixed modifications to carbamidomethylation of Cys. MSBooster, Percolator and ProteinProphet default options were used, and results were filtered by 1% FDR at protein level. The searched results were then imported to Scaffold 5 Q+S (Proteome Software Inc., v5.3) for data visualization.

Data from the second experiment (pull-down of Rv3035-3xFLAG) was acquired using a NanoAcquity UPLC (Waters Corporation, Milford, MA) coupled to an Orbitrap Fusion Lumos Tribrid (Thermo Fisher Scientific, Waltham, MA) mass spectrometer. Peptides were trapped and separated using an in-house packed precolumn packed with 2cm ReproSil-Pur C18-AQ (Dr. Maisch, GmbH, 120 Å, 5 μm) particles plus and an in-house 75 μm I.D. fused silica analytical column (as previously described) and the same solvent composition. 3.8µL sample was injected, and the peptides were trapped at a flow rate of 4µL/min with 5% mobile phase B for 4 min, followed by a gradient elution with 5-35% B at a flow rate of 300nL/min over 120 min (total run time 145 min). Mass spectra were acquired over m/z 375-1500 Da with a resolution of 120,000 (m/z 200). Tandem mass spectra were acquired using data-dependent acquisition with an isolation width of 1.6 Da, HCD collision energy of 30%, resolution of 15,000 (m/z 200), maximum injection time of 50 ms, and an AGC target of 50,000.

Raw data files from the Orbitrap Fusion Lumos Tribrid were peak processed with Proteome Discoverer (version 2.5, Thermo Scientific) followed by identification using Mascot Server (Matrix Science, v2.6.2) against the same database used in the previous experiment. Search parameters were as follows: tryptic digestion with up to 2 missed cleavages; peptide N-terminal acetylation, methionine oxidation, and N-terminal glutamine to pyroglutamate conversion were specified as variable modifications. Carbamidomethylation of cysteines was set as static modification. Assignments were made using a 10-ppm mass tolerance for the precursor and 0.6 Da mass tolerance for the fragment ions. All non-filtered search results were processed by Scaffold 5 Q+S at 1.0% false-discovery rate (FDR) for peptides and 99% threshold for proteins (2 peptides minimum).

For each experiment, Total Spectrum Count (TSC) was exported from Scaffold 5 Q+S into a Microsoft Excel spreadsheet. Zeros were replaced with ones in WT control samples to allow fold-change (FC) calculation of every protein detected in the samples containing the bait protein. For normalization, we summed the TSC values for all proteins in each replicate, calculated the average of the sum of the three replicates and divided the average by the sum value of each replicate (this is the normalization factor). Then, we multiplied each protein TSC by the normalization factor of that replicate sample, calculated the median, log2FC of tagged group over WT control, and p-value by performing T-tests of the normalized values in each row. P-values were transformed into -log10(p-value) for visualization of the data as a volcano plot.

### Co-essentiality analysis

We performed co-essentiality analysis on CRISPR interference (CRISPRi) data (*13*), which profiled Mtb fitness under 80 experimental conditions. These conditions span 9 antibiotics (BDQ = bedaquiline; CLR = clarithromycin; EMB = ethambutol; INH = isoniazid; LVX = levofloxacin; LZD = linezolid; RIF = rifampicin, STR = streptomycin; VAN = vancomycin), each tested under 9 screening conditions defined by a combination of pre-depletion time (1, 5, or 10 days of CRISPRi induction before drug exposure) and drug concentration (“Low,” “Medium,” or “High” partially inhibitory doses). We used only the log₂ fold change (LFC) values from negative selection screens, which report the fitness effects of gene knockdown.

Each gene was thus represented by an 80-dimensional LFC vector capturing its essentiality profile across the drug/time/concentration matrix. To account for covariance across screens— such as batch effects—we computed an 80 × 80 empirical covariance matrix (Σ) from the LFC data. We then applied generalized least squares (GLS), following a previously reported approach (*53*) to compute pairwise co-essentiality relationships among 3,979 genes. This yielded a 3,979 × 3,979 matrix of GLS coefficients and corresponding p-values.

We corrected for multiple hypothesis testing using the Benjamini–Hochberg procedure (fdrcorrection in the Python statsmodels package). Gene pairs with a false discovery rate (FDR)- adjusted p-value < 0.001 were considered significantly co-essential, resulting in approximately 16,000 high-confidence gene pairs.

### Design of expression constructs for *M. thermoresistibile* (Mth) Rv3035

Initial attempts to express Mtb Rv3035 in *E. coli* using N- or C-terminal 6xHis-tag (His) constructs with full-length and various N-terminal truncations failed to produce soluble Rv3035. To improve solubility, we turned to a thermostable homolog of Rv3035, Mth Rv3035, Accession no. WP_040546537, and constructed two *E. coli* codon-optimized constructs cloned into pET28a with a TEV-cleavable N-terminal His-tag (GenScript, Piscataway, NJ). The Mth Rv3035 construct that encodes for the mature form of Rv3035 without its predicted signal peptide (SignalP 6.0 (*14*)), starting at residue Gly2 (fig. S4B), produced soluble Rv3035 only when expressed in *E. coli* T7SHuffle cells (New England Biolabs, Ipswich, MA). Notably, T7SHuffle cells promote disulfide bond formation (*54*), and Rv3035 is predicted to harbor two disulfide bonds.

### Generation of FecB mutants for *in vitro* studies

A pET28a FecB construct encoding Mtb FecB (without the signal peptide, mature FecB starting at residue 39) with a thrombin-cleavable N-terminal 6xHis-tag (*7*) was used as the template for *in vitro* site-directed mutagenesis using Pfu Ultra Fusion HS DNA polymerase (Agilent) with the primers listed in table S5 and then confirmed by DNA sequencing (GeneWiz from Azenta Life Sciences). The FecB construct encoding no His-tag was generated using the pET28a FecB plasmid and excising the encoded *fecB* gene with *Nde*I and *BamH*I and ligating it into a pET22b vector.

### Expression and purification of Mth Rv3035 and the Mth Rv3035-Mtb FecB complex

Mth Rv3035-His was expressed in *E. coli* T7SHuffle cells in Terrific Broth (TB) media containing 0.4% glycerol and 50μg/mL kanamycin (Kan). To express the Mth Rv3035-Mtb FecB complex, which we refer to as the FecB-Rv3035 complex throughout the manuscript, both pET28a Mth Rv3035-His and pET22b Mtb FecB plasmids were co-transformed into T7SHuffle cells with selection on LB + 50μg /mL Kan and 100μg/mL ampicillin (Amp) agar plates. A fresh transformant was used to inoculate TB containing 0.4% glycerol, 50μg/mL Kan and 100μg/mL Amp. Cells were grown at 37°C to an OD_600nm_ of ∼ 0.6, before protein expression was induced by the addition of 0.5mM isopropyl β-D-1-thiogalactopyranoside (IPTG). The cells grown overnight at 20°C and harvested by centrifugation.

Both Mth Rv3035-His and the FecB-Rv3035-His complex were purified using the same protocol. Cell pellets were resuspended in 10x vol(mL)/w(g)of cells in Buffer A (150mM NaCl, 50mM Tris, pH 7.8) with 0.25mM phenylmethylsulfonyl fluoride (PMSF) with stirring at 4°C for 30 minutes. Cells were lysed by sonication and cell debris was pelleted by centrifugation at 14,000rpm for 1h at 4°C. The cell lysate was passed through a 1µm syringe filter to remove insoluble debris and applied to a His-trap column (Cytiva, Marlborough, MA) pre-equilibrated with Buffer A using an AKTA Start FPLC instrument (Cytiva, Marlborough, MA). Following a 10x column volume (CV) wash step with Buffer A, a gradient of Buffer A from 0 to 500mM imidazole over 20 CVs elute His-tagged proteins. Fractions with containing Mth Rv3035-His or the FecB-Rv3035-His complex were concentrated using an Amicon concentrator (MilliporeSigma, 30kDa cut-off) with buffer exchange into Buffer A.

### Expression and purification of Mtb FecB and mutants

Purification of WT FecB and FecB mutant proteins were carried out in a similar manner as previously described (*7*). FecB WT or mutants were expressed in BL21-(DE3) cells and cells were grown in TB + 0.4% glycerol + 50μg/mL Kan at 37°C to an OD_600nm_ of ∼0.6, before protein expression was induced with IPTG (0.5mM). Cell growth was continued overnight for 20h at 20°C, before cells were harvested by centrifugation. Cell pellets were resuspended in 10x vol(mL)/w(g) Buffer B (350 mM NaCl, 50 mM Tris, pH 7.8, 10% glycerol) with 0.25mM PMSF with stirring at 4°C for 30 minutes. Cells were lysed by sonication and cell debris was pelleted by centrifugation at 14,000rpm for 1h at 4°C. The cell lysate was filtered and applied to a Ni-NTA resin pre-equilibrated with Buffer B using gravity flow. The column was washed with Buffer B and eluted with a stepwise elution gradient of imidazole going up to 600mM imidazole in Buffer B. Fractions found to contain FecB by SDS-PAGE analysis (expected MW ∼37 kDa) were pooled and dialyzed against Buffer B overnight at 4°C.

### Removal of N-terminal His-tag from Rv3035-His and FecB-His

For cleavage of His-tags from Rv3035-His and from the FecB-Rv3035-His complex, TEV was added to approximately 30-100mg of protein at a final concentration ∼0.1mg/ml TEV and incubated overnight at 4°C. The cleavage reaction was passed over a Ni-NTA column pre-equilibrated with Buffer A + 10mM imidazole and chased with Buffer A + 10mM imidazole. Flow-through fractions containing Rv3035 or FecB-Rv3035 complex were concentrated and buffer exchanged into Buffer A.

For the isolation of tag-free WT FecB, which was used for some ITC experiments, a similar procedure was used but with thrombin at ∼30U thrombin:1mg FecB-His. The cleavage reaction was passed over a Ni-NTA column pre-equilibrated with Buffer B + 10mM imidazole and chased with Buffer B + 10mM imidazole. Flow-through fractions containing WT FecB were concentrated and buffer exchanged into Buffer B.

### Isothermal titration calorimetry (ITC) experiments

ITC experiments were carried out using a Malvern MicroCal PEAQ-ITC instrument (Malvern, United Kingdom). Prior to the experiments, proteins were dialyzed into Buffer B and the same dialysis buffer was used for protein dilutions. 19-point titrations were carried out using 200μM Mth R3035 in the syringe and 20μM FecB in the cell, whereas 13-point titrations were carried out using 300μM Rv3035 and 30μM FecB-His mutant protein, or 200μM Rv3035-His and 20μM FecB-His. Notably, the His-tag on FecB did not appreciably change the affinity of FecB for Rv3035. Curve fitting was carried out using the Malvern instrument analysis software.

### Structure determination of Mth Rv3035 and the FecB-Rv3035 complex

Initial screening of Mth Rv3035 (15mg/mL) and FecB-Rv3035 complex (38mg/mL) were carried out in 96-well trays using a Mosquito liquid handler (SPT Labtech, Cambridge, MA) and commercially available crystallization screens by the hanging drop vapor diffusion method.

#### Mth Rv3035 alone

Diffraction-quality Mth Rv3035 crystals were grown with a crystallization reservoir of 0.1M sodium chloride; 0.1M HEPES, pH 7.5; 1.6M ammonium sulfate with Rv3035 at 15 mg/mL and a reservoir to protein ratio of 1:1 at room temperature. The crystals were cryocooled in mother liquor supplemented with 25% glycerol. Data was collected at SSRL 12-2 at the wavelength 1 Å. Data was processed in xds (*55*) to a resolution of 1.9 Å (table S1). A model of Mth Rv3035 was generated on the AlphaFold server (*26*) and was used for molecular replacement (MR) in phaser (*56*) in phenix. The MR solution placed two subunits in the asymmetric unit (ASU) and was improved by autobuild in phenix (*57*), and refined in phenix.refine(*58*). Following autobuild, the N-terminal residues 10-37 were built *de novo* in coot (*59*). The N-terminal 9 residues of Rv3035 could not be modeled for both subunits, and for subunit B, the last C-terminal residue also could not be modeled due to unresolved electron density. The model was further refined in phenix.refine and coot to a final R_work_/R_free_ of 16.1 / 20.25 (table S1).

#### FecB-Rv3035 complex

Diffraction-quality crystals of the FecB-Rv3035 complex were obtained with a crystallization reservoir of 3M sodium chloride; 0.1M Bis-Tris, pH 7.0 with a reservoir to protein complex ratio of 1: 1. To further improve diffraction quality of the FecB-Rv3035 complex, an additive screen was performed (Hampton Research, Aliso Viejo, CA), with the final conditions used being a crystallization reservoir of 3M sodium chloride; 0.1M Bis-Tris, pH 7.0; 3% trehalose, with the Rv3035-FecB complex at 38mg/ml and a reservoir to protein ratio of 1:1. Crystals were cryoprotected in reservoir solution with 25% glycerol and flash frozen with liquid nitrogen. Data was collected at SSRL 12-2 at the wavelength 1 Å.

Data was processed in XDS (*55*) to a resolution of 3.3 Å (table S1). The phases were solved by MR in phaser (*56*) in phenix, placing a single subunit of the deposited FecB structure (PDB ID 7UQ0,(*7*)) in the ASU in complex with a single subunit of the Mth Rv3035 structure solved above. The MR solution was improved in autobuild and refined as described above in phenix.refine (*58*) and coot (*59*) to a final R_work_/R_free_ of 25.2/26.3 (table S1).

### Knockout construction

Δ*rv3035* and Δ*fecB*Δ*rv3035* mutants were generated using a suicide plasmid approach as previously described (*60*) with a few modifications. Constructs containing 500bp fragments corresponding to regions upstream and downstream of *rv3035* were ordered from GenScript and cloned into a temperature-sensitive deletion vector pDE43-XSTS to flank the zeocin resistance cassette and generate pKO-XSTS-Zeo-*rv3035* via Gateway Cloning. Mtb WT and Δ*fecB* were transformed with both the pKO-XSTS-Zeo-*rv3035* and a *rv3035* complementation plasmid (pGMCS-T02-P*hsp60*-SD1-*rv3035*_A-HA) to generate a merodiploid strain since initial attempts of generating the knockout with pKO-XSTS-Zeo-*rv3035* alone had failed. Transformants were plated on Middlebrook 7H10 agar (BD #262710) containing 0.5% glycerol, 10% OADC (BD #212351) and 25μg/mL Zeo (for WT) or 50μg/mL Hyg and 25μg/mL Zeo (for Δ*fecB*) and incubated at 37°C until colonies were approx. 1mm in diameter. Colonies were tested for reporter gene XylE activity directly on plates by addition of a drop of catechol solution (1g in 20mL water). After 30 minutes, candidate colonies that turned yellow were selected and grown in regular 7H9 with appropriate antibiotics until they reached an OD_580nm_ of 1-1.5. Candidates were then grown on 7H10 agar with 10% sucrose at 37°C until colonies were approx. 1mm in diameter. Colonies were test for reporter gene activity as above, but this time white colonies (that had successfully lost the suicide plasmid) were selected. Genomic DNA (gDNA) was extracted from a few mutant candidates and allelic exchange was confirmed by PCR using primers listed in table S6. To obtain the final knockout mutants, we replaced the *rv3035* complementation plasmid integrated into attL5 with an empty plasmid containing an apramycin resistance cassette (pGMCA-OXOX). Transformants were plated on 7H10 containing Zeo (or Zeo and Hyg) and 60μg/mL apramycin. Δ*rv3035* and Δ*fecB*Δ*rv3035* mutants took about 6-8 weeks to form visible colonies. Candidate mutants were selected and tested for streptomycin sensitivity in liquid culture as an indication that the copy of *rv3035* (carrying streptomycin resistance) was successfully replaced in attL5. Finally, gDNA was extracted for whole genome sequencing to confirm deletion of *rv3035* and insertion of zeocin and apramycin cassettes.

### Cell envelope permeability assays

BODIPY Fluorescent (FL) vancomycin (Invitrogen #V34850) uptake was performed as previously reported (*5*) with a few modifications. Briefly, mid-log phase Mtb cultures were washed once and adjusted to an OD_580nm_ of 2 in PBS and incubated with 2μg/mL of BODIPY FL vancomycin. 350μL sample aliquots were taken at the 30 minutes incubation time point, washed twice in PBS 0.05% Tween80, and resuspended in 350μL PBS. Negative control samples were treated the same way but did not receive BODIPY FL vancomycin. 100μL of each sample was distributed in triplicate wells in a black-sided 96-well plate. Fluorescence was measured (excitation wavelength of 485nm and emission wavelength of 538nm) and normalized to the OD_580nm_ of the final suspension to account for cell loss during washing steps.

For the calcein-AM (Invitrogen #C3099) uptake assay, mid-log phase cultures were adjusted to an OD_580nm_ of 0.8 in regular 7H9 and 100μL aliquots were added to the wells of a black-sided 96-well plate in triplicates. 100μL of calcein-AM (1μg/mL in 7H9 0.4% glucose) was added to the wells resulting in a final concentration of 0.5µg/mL calcein-AM. Fluorescence was measured (excitation wavelength of 495nm and emission wavelength of 520nm) at 2 minutes intervals over a course of 60 minutes. Fluorescence was normalized to OD_580nm_ measured at 60 minutes.

### Spot assay

Mig-log phase Mtb cultures were adjusted to an OD_580nm_ of 0.3 and 10-fold serial dilutions in PBS 0.05% Tween80 were spotted on 7H10 agar with 5µL per spot. Plates were incubated at 37°C and imaged after 2-3 weeks. To confirm colony morphology alteration and growth defect of Δ*fecB*, Δ*rv3035* and Δ*fecB* Δ*rv3035* mutants, plates were incubated for 4-5 additional weeks, after which the phenotypes were unchanged.

### DMN-Tre labeling and flow cytometry

Mid-log phase Mtb cultures were adjusted to OD_580nm_ of 0.5 in regular 7H9. For CRISPRi strains, cultures were split into two flasks and 200ng/mL of ATc was added to one of the flasks 2-3 days prior to labeling with DMN-Tre to pre-deplete the gene of interest. DMN-Tre probe (*31*) was added to 500µL of culture at a final concentration of 100µM. Samples were incubated at 37°C overnight for approximately 20h. On the next day, cultures were spun down at 10,000rpm for 3 minutes, and the supernatant containing DMN-Tre was discarded. Samples were fixed with 4% PFA (Biolegend #420801) for 4h at room temperature. After two washes in PBS 0.05% Tween80, bacteria were resuspended in PBS. Data collection was performed on a FACSymphony A5 Cell Analyzer (BD Biosciences) at the Weill Cornell Medicine Flow Cytometry Core Facility using the 405nm violet laser and the 405 [425/50] detector. Fluorescence data were obtained for 200,000 events per sample and analyzed in FlowJo software (BD) by first gating on the bacterial population in the side scatter versus forward scatter plot followed by calculation of the mean fluorescence intensity.

### Preparation and analysis of lipoglycans and arabinogalactan

Lipoglycan and mycolyl-arabinogalactan-peptidoglycan (mAGP) complex extraction from delipidated cells followed earlier procedures (*61*, *62*). Determination of the glycosyl linkage patterns of lipoglycans and mAGP followed earlier procedures (*61*). Per-*O*-methylated alditol acetates were analyzed by GC/MS on a Thermo Scientific TRACE 1310 Gas Chromatograph paired with a Thermo Scientific TSQ 8000 Evo Triple Quadrupole GC-MS/MS. Samples were run on a Zebron ZB-5HT Inferno 30 m x 0.25 mm x 0.25 µm capillary column (Phenomenex) at an initial temperature of 100°C. The temperature was increased to 150°C at a ramp rate of 20°C min^-1^, then to 240°C at a ramp rate of 5°C min^-1^ and was held at this temperature for 3 min to be finally increased to 300°C at a rate of 30°C min^-1^ and held at the final temperature for 5 min. Data handling was carried out using the Thermo Scientific Chromeleon Chromatography Data System software.

### Mouse infection

Sixty-four female, eight-week-old *Mus musculus* C57BL/6 (Jackson Laboratories) were infected with ∼100 colony forming units (CFU)/mouse using an Inhalation Exposure System (Glas-Col). Mice in the doxycycline group started to receive doxycycline chow 2 days before aerosol infection. The FecB-TetON strain was grown to mid-log phase in the presence of 1μg/mL ATc, a single cell suspension was prepared in PBS 0.05% Tween80 and then resuspended in PBS for a final OD_580nm_ of 0.2. Lungs and spleen were homogenized in PBS and plated on 7H10 agar containing 1μg/mL ATc to determine CFU/organ at the indicated time points. This experiment was performed in accordance with the Guide for the Care and Use of Laboratory Animals of the National Institutes of Health, with approval from the Institutional Animal Care and Use Committee of Weill Cornell Medicine.

### Quantification and statistical analysis

Data analysis and generation of graphs were performed in Prism version 10.5 (GraphPad).

## Acknowledgements

We thank Sae Woong Park (Weill Cornell Medicine) for sharing the Δ*mbtK* and Δ*mbtI* strains, Carolina Trujillo (Weill Cornell Medicine) for assistance with the animal experiment, and Allison Fay (Memorial Sloan Kettering Cancer Center) for her advice on the thin layer chromatography experiment. We thank the staff of the University of Massachusetts Chan Medical School Mass Spectrometry Facility and Kadamba Papavinasasundaram for their help with the immunoprecipitation tandem mass spectrometry experiments. We thank the Kamariza Lab at the University of California, Los Angeles for gifting an aliquot of the DMN-Tre probe. We thank the staff of the Weill Cornell Medicine Genomics Resources Core Facility and Flow Cytometry Core Facility for their contribution and advice on the RNA-seq and flow cytometry experiments, respectively. We thank the staff of the Advanced Light Source at Berkeley National Laboratories and the Stanford Synchrotron Radiation Lightsource for their invaluable help in data collection. We thank the Analytical Resources Core Facility at CSU (RRID: SCR_021758) for its help with GC/MS analyses.

## Funding

This work was supported by the National Science Foundation Graduate Research Fellowship under grant No. 2139291 and a Potts Memorial Foundation Pre-Doctoral Fellowship (to T.K.). This work was also supported by the National Institute of Allergy and Infectious Diseases (NIAID)/ National Institutes of Health (NIH) grant AI155674 (to M.J.), grant K08AI148584 (to C.B.) and grant AI095208 (to C.W.G.). Additionally, this work was funded by the NIAID/NIH predoctoral training grant support AI141346 (to R.d.M. and J.M.). The content is solely the responsibility of the authors and does not necessarily represent the official views of the NIH.

## Author contributions

Conceptualization: T.K., D.S. and S.E. Methodology: T.K., D.S. and S.E. Visualization: T.K. and S.E. Investigation: T.K., C.B., C.D.H., B.J.C., A.S., A.J., L.C., S.K.A., J.M., J.M., R.d.M. and H.K. Formal analysis: T.K. and S.E. Project administration: T.K. and S.E. Validation: T.K. Supervision: M.J., K.R., C.W.G. and S.E. Funding acquisition: T.K., M.J., C.B., C.W.G., R.d.M., J.M. and S.E. Writing—Original draft: T.K., C.D.H., B.J.C., M.J., C.W.G. and S.E. Writing—Review and editing: T.K., C.D.H., B.J.C., A.J., D.S., M.J., C.W.G. and S.E.

## Competing interests

The authors declare that they have no competing interests.

## Data and materials availability

All data needed to evaluate the conclusions in the paper are present in the paper and/or the Supplementary Materials. Bacterial strains generated in this paper will be provided upon request pending scientific review and a completed material transfer agreement. Requests should be directed to S.E. at.

## Notes

### Competing Interest Statement

The authors have declared no competing interest.

### Summary of Updates

We added new data that show the analysis of lipoglycans and arabinogalactan in wild type and mutant Mtb strains. This led to three new authors from Colorado State University.

