## Supplementary Materials for "A periplasmic protein complex supports arabinofuranosyltransferase activity and mediates intrinsic drug resistance in *Mycobacterium tuberculosis*"

This file contains supplementary materials and methods, figures S1-14, tables S1-6 and description of datasets S1-5.

### Supplementary materials and methods

#### Western blot analysis

Mtb strains were grown to $OD_{580nm}$ of ~1, pelleted at 4000rpm for 8 minutes and washed once with PBS. Pellets were resuspended in 600μL of PBS with protease inhibitor (cOmplete Mini EDTA-free Protease Inhibitor Cocktail, Roche), transferred to 2mL screw cap bead-beater tubes (Labcon # 4-3661-875-000), and bead-beaten with 0.1mm zirconia/silica beads (Biospec #11079101z) in a Precellys Evolution Touch homogenizer (Bertin technologies) 4x for 30 seconds at 7,500rpm, with a 30 second incubation on a cooling rack in between each bead-beating. The tubes were then centrifuged once at 8,000 x g for 30 seconds at 4°C to separate the zirconia beads from the lysate. The bead-free supernatant was then centrifuged at 11,000 x g for 5 minutes at 4°C to remove cell debris. This whole-cell lysate was mixed with 4x Laemmli sample buffer (Bio-Rad #1610747) containing β-mercaptoethanol at a 1:4 dilution and boiled at 95°C for 10 minutes.

For SDS-PAGE analysis, 10-20μL of each sample was loaded onto a 4-20% Bio-Rad PROTEAN TGX pre-cast gel and run at 120V for 1h in 1x Tris-glycine buffer. The Precision Plus Protein Dual Xtra Prestained Protein Standards (Bio-Rad #1610377) was used as a molecular weight reference. Using the semi-dry iBLOT (Invitrogen) transfer method, the gels were transferred onto nitrocellulose membranes. Membranes were blocked with LICOR intercept PBS blocking buffer (LICOR #927-70001) for 1h at room temperature, followed by overnight incubation with the primary antibody in antibody diluent (1:1 blocking buffer: PBS with 0.1% (v/v) Tween20) at 4°C. After incubation with the primary antibody, membranes were washed with PBS 0.1% Tween20 3x for 6 minutes and then incubated with secondary antibody (diluted in antibody diluent solution) for 1-2h at room temperature. After washing with PBS 0.1% Tween20 3x for 6 minutes, the blots were imaged using the Azure 600 bio-imager. The anti-FecB antisera was obtained from rabbits 58 days after exposure to purified recombinant FecB (ThermoFisher AB3448, D58) and used at a 1:5,000 dilution. The rabbit anti-PrcB antibody (GN-13.782) was diluted 1:10,000. The mouse anti-FLAG antibody (Sigma #F3165,) was diluted 1:800. The mouse anti-HA antibody (Sigma #H3663) was diluted 1:10,000. Secondary antibodies IRDye® 800 donkey anti-rabbit IgG antibody (H + L) (LICOR #926-32213) was diluted 1:15,000, IRDye® 680 donkey anti-rabbit IgG antibody IgG (H + L) (LICOR #926-68073) was diluted 1:10,000, and IRDye® 800CW goat anti-mouse IgG antibody (H + L) (LICOR #926-32210) was diluted 1:10,000.

**Fluorescence titrations for FecB mutants with apo-carboxymycobactin**

Fluorescence quenching titrations of apo-carboxymycobactin (cMB) were performed as described previously (*63*, *64*). Stock solutions of FecB (100 nM) and cMB (80 μM) were prepared in 50 mM Tris-HCl pH 7.4, 150 mM NaCl. cMB was titrated into FecB and in between each titration the solution was incubated for 3 minutes with stirring at 200 rpm at 20 °C. Fluorescence spectra were acquired between 300 – 500 nm using a Hitachi F-7100 Fluorescence Spectrophotometer through excitation at 285 nm with the following settings: PMT voltage of 950 V, excitation slit width of 2.5 nm, emission slit width of 5.0 nm, and a scan speed of 240 nm/min.

Results from the fluorescence-based assay were fit to Eqn. 1 (*64*) to determine the equilibrium dissociation-constant ($K_D$) of cMB with FecB and its mutants.

*Eqn. 1*:

$$F = \frac{([FecB] + [cMB] + K_d) - \sqrt{([FecB] + [cMB] + K_d)^2 - 4[FecB][cMB]}}{2} \times \left(\frac{F_{min} - F_{max}}{[FecB]}\right) + F_{max}$$

In Eqn. 1, [FecB] is the total concentration of FecB or mutants, [cMB] is the total concentration of cMB, $F_{\mathrm{max}}$ is the emission intensity without ligand, and $F_{\mathrm{min}}$ is the emission intensity for fully cMB-bound FecB (2). Fitting of the fluorescence emission intensity at 335 nm for $K_D$ determination was performed using GraphPad Prism (Ver 9.3.1).

**Size-exclusion chromatography**

Samples of Mth Rv3035-His, the FecB-Rv3035-His complex, and Mtb FecB-His were individually applied to a Superdex 200 10/300 column (Cytiva, Marlborough, MA) in column in Buffer A (150mM NaCl, 50mM Tris, pH 7.8) using an AKTA FPLC instrument (Cytiva, Marlborough, MA). Apparent molecular weights were calculated based on protein standards run on the same column in the same buffer on the same day the test proteins were run.

**Circular dichroism (CD) of Mtb FecB and its mutants**

All experiments were performed at 25°C using a Jasco J-180 spectropolarimeter with a Digital Integration Time of 1 second and a bandwidth of 1 nm. Sample wavelengths were read from 260-190nm at 100 nm/min for a total of 10 accumulations. Sample preparation involved a 10x dilution in water using a 10mm path-length quartz cuvette. The BeStSel tool (https://bestsel.elte.hu/index.php) was used to quantify secondary structural elements.

**Protein interaction nickel-affinity pull-down assays**

A mixture of 3μM Mth Rv3035 and Mtb FecB-His (WT or mutants) were incubated with 100μL Ni-NTA resin in 200μL Buffer B (350 mM NaCl, 50 mM Tris, pH 7.8, 10% glycerol) + 10mM imidazole for 30 minutes at room temperature followed by a short low-speed spin to pellet the resin, where supernatant was the flow-through. Next, 200µL Buffer B + 10mM imidazole was used to resuspend the resin followed by a short low-speed spin; the supernatant was the wash. As a low imidazole elution step, 200µL Buffer B + 50mM imidazole was used to resuspend the resin followed by a brief low-speed spin, where the supernatant was the low imidazole elution. As a final elution step, 200µL Buffer B with 400mM imidazole was used to resuspend the resin followed by a brief low-speed spin, and the supernatant was the high imidazole elution. The input and resulting supernatants were run on an SDS-PAGE gel.

**Co-immunoprecipitation assay**

Similarly to pull-down tandem mass spectrometry, whole-cell lysates were collected from log-phase cultures grown in the shaking incubator and incubated with 1% n-dodecyl -D-maltoside for 2h on a rotator at 4°C. Next, samples were incubated with anti-DYKDDDDK (FLAG) magnetic agarose (Pierce #A36797) or anti-HA magnetic beads (Pierce #88836). The flow through from the magnetic agarose/bead incubation was saved for downstream analysis. Captured proteins were dissociated from magnetic agarose/bead by addition of 3x DYKDDDDK (3xFLAG) peptide (Pierce #A36805) or HA peptide (Pierce #26184), according to the manufacturer's instructions. 4x Laemmli sample buffer (Bio-Rad #1610747) containing β-mercaptoethanol was added to whole-cell lysate, flow through and peptide eluate fractions at 1:4 dilution. PBS containing 4x Laemmli buffer was added to beads and all fractions were boiled at 95°C for 10 minutes. The supernatant collected from the boiled beads after magnetic separation is referred to as the boiled beads eluate.

**AlphaFold 3 Complex Predictions**

Mature versions of FecB and Rv3035 without the signal peptide sequence (FecB starting at residue 39 and Rv3035 starting at residue 28) were used in all multimer complex predictions. AlphaFold 3 (*26*) was used to model different combinations of FecB, Rv3035, AftB and Rv0227c. All data shown in fig. S11 and S12 were obtained from the AlphaFold 3 website (https://alphafoldserver.com/).

**Thin-Layer Chromatography**

Approximately 300µg of lipids extracted from the supernatant of AOT-treated samples (referred to as outer leaflet of the mycomembrane, or OM) was spotted on high-performance thin layer chromatography (HPTLC) plates (Miles Scientific #P60077), run 3 times in chloroform-methanol-water (90:10:1) and allowed to air dry. HPTLC plates were visualized by spraying with 5% molybdophosphoric acid in ethanol and charring. 10µg of the trehalose dimycolate (TDM) standard (Sigma #T3034) was run with the lipid samples in each HPTLC plate. Quantification of TDM band intensity (densitometry analysis) was performed in Fiji (*65*) as previously described (*66*) with a few modifications. Steps 2-10 of the "Western Blot and Protein Band Quantification" section (*66*) were followed and the area of the peak within each lane was extracted for statistical analysis.

### Supplementary figures

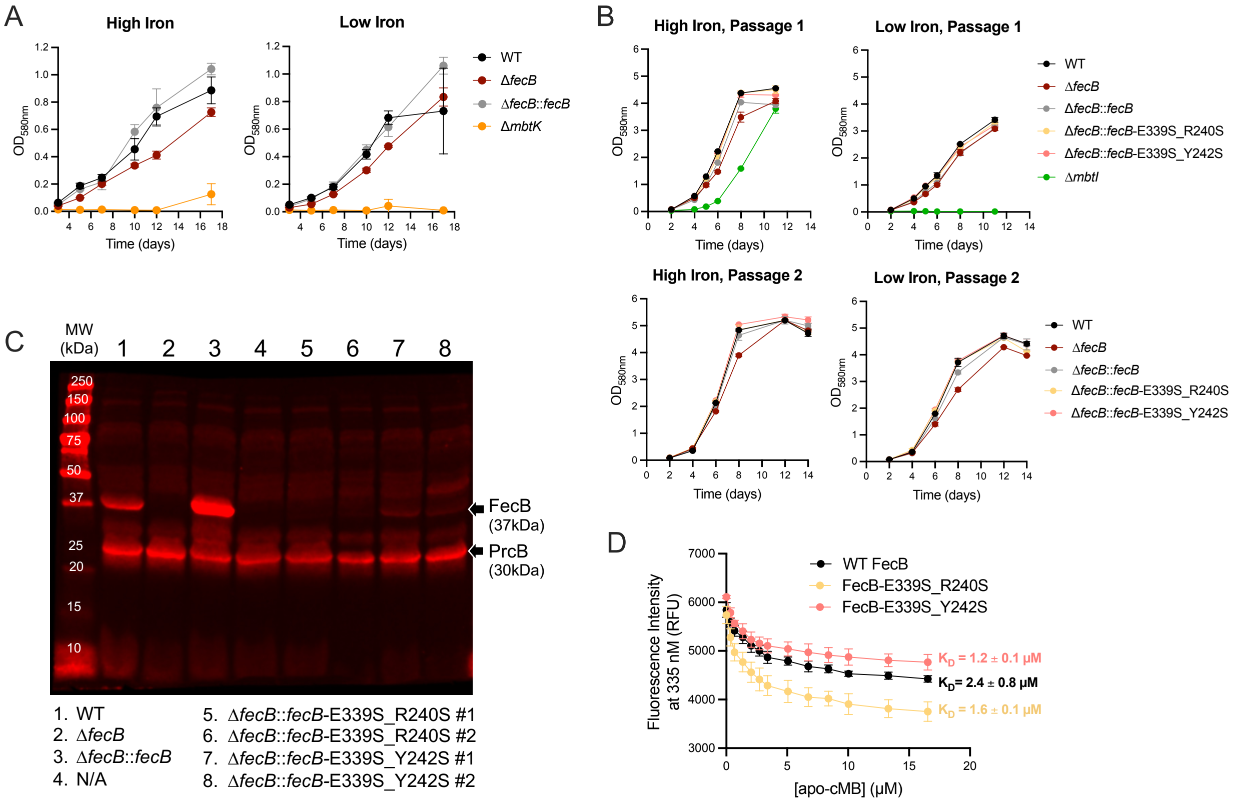


**Fig. S1. Growth curves in iron-limiting conditions and characterization of FecB E339S_R240S and E339S_Y242S mutants.** (A) Growth curves of iron pre-depleted strains in iron-chelated detergent-free media with 160μM (high iron) or 1μM (low iron) AFC. Δ*mbtK* (mycobactin biosynthesis mutant) was used as a positive control that cannot grow in iron-limiting conditions. (B) Growth curves of iron pre-depleted strains in iron-chelated media with 160μM (high iron) or 1μM (low iron) AFC. Cultures in low iron were grown until day 11 (Passage 1) and diluted to starting OD=0.025 in high/low iron media (Passage 2). Δ*mbtI* (mycobactin biosynthesis mutant) was used as a positive control that cannot grow in iron-limiting conditions. (C) Western-blot analysis of whole cell lysates, #1-2 after strain name indicate individual colonies (biological replicates). Nitrocellulose membrane was probed with anti-FecB antiserum and anti-PrcB antiserum (loading control). (D) Representative fluorescent emission intensities at 335nm after excitation at 280nm of 100nM WT FecB and FecB variants E339S_R240S and E339S_Y242S with increasing concentrations of apo-cMB. Curves were fit using a previously published equation (*67*) and apo-cMB affinities ($K_D$) are included for each titration. Error bars: ±SD.

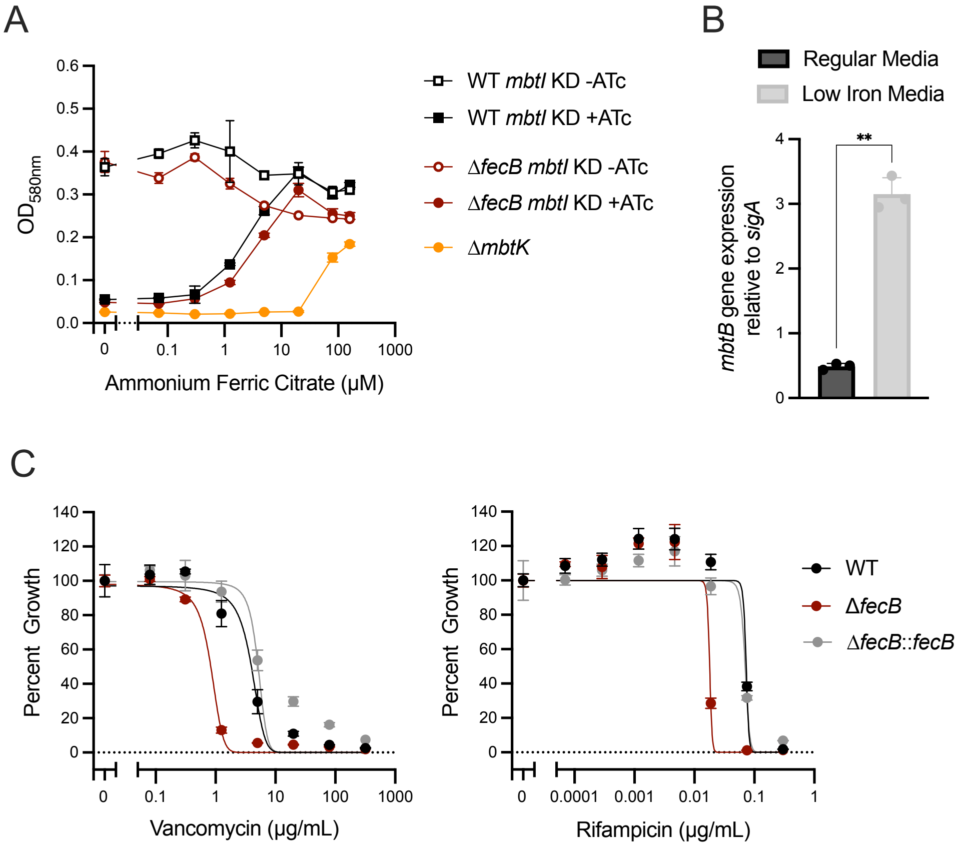


**Fig. S2. Validation of *mbtI* KD strains, *mbtB* induction in low iron media and minimal inhibitory concentration assay in detergent-free conditions.** (A) Growth curve of iron pre-depleted strains in iron-chelated detergent-free media with increasing concentrations of AFC, in the absence or presence of CRISPRi KD inducer ATc, day 7. Δ*mbtK* (mycobactin biosynthesis mutant) was used as a positive control that cannot grow in iron-limiting conditions. (B) Differential expression by reverse transcription quantitative PCR (RT qPCR) of representative siderophore biosynthesis gene *mbtB* in WT strain grown in regular media and low iron media (iron-chelated media with 1μM AFC). Statistical significance was determined by unpaired t-test with Welch's correction. (C) MIC assays for vancomycin and rifampicin in detergent-free media, percent growth calculated from no drug control wells, day 14. Experiments in A and C were performed in 96-well plates with wells resuspended prior to $OD_{580nm}$ measurement. Error bars: ±SD. **P* < 0.05; ***P* < 0.01; ****P* < 0.001; *****P* <0.0001.

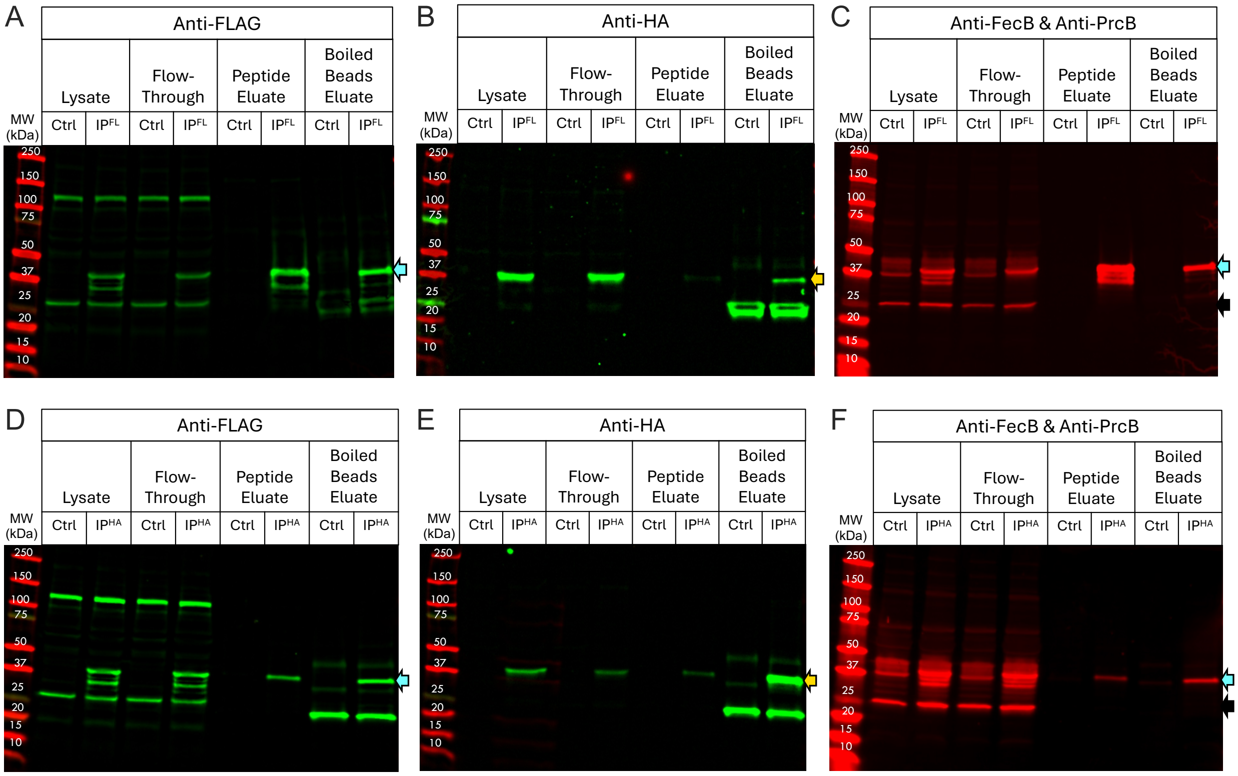


**Fig. S3. Co-immunoprecipitation of FecB-FLAG and Rv3035-HA.** Western blot analysis of protein fractions from Δ*fecB* harboring plasmids encoding FecB-FLAG and Rv3035-HA immunoprecipitated with FLAG (A-C, $IP^{FL}$) or HA (D-F, $IP^{HA}$) magnetic beads. WT lysate without tagged proteins was used as a control (Ctrl). Nitrocellulose membranes were probed with anti-FLAG primary antibody (A, D) to detect FecB (~37kDa, blue arrows), anti-HA primary antibody (B, E) to detect Rv3035 (~37kDa, truncated version with 360 amino acids, yellow arrows), and anti-FecB antiserum (C, F) to detect FecB (~37kDa, blue arrows). Anti-PrcB primary antibody (C, F) was used to detect PrcB (~30kDa, observed at lower molecular weight, black arrows) as a loading control. MW = Molecular Weight, kDa = kilodalton.

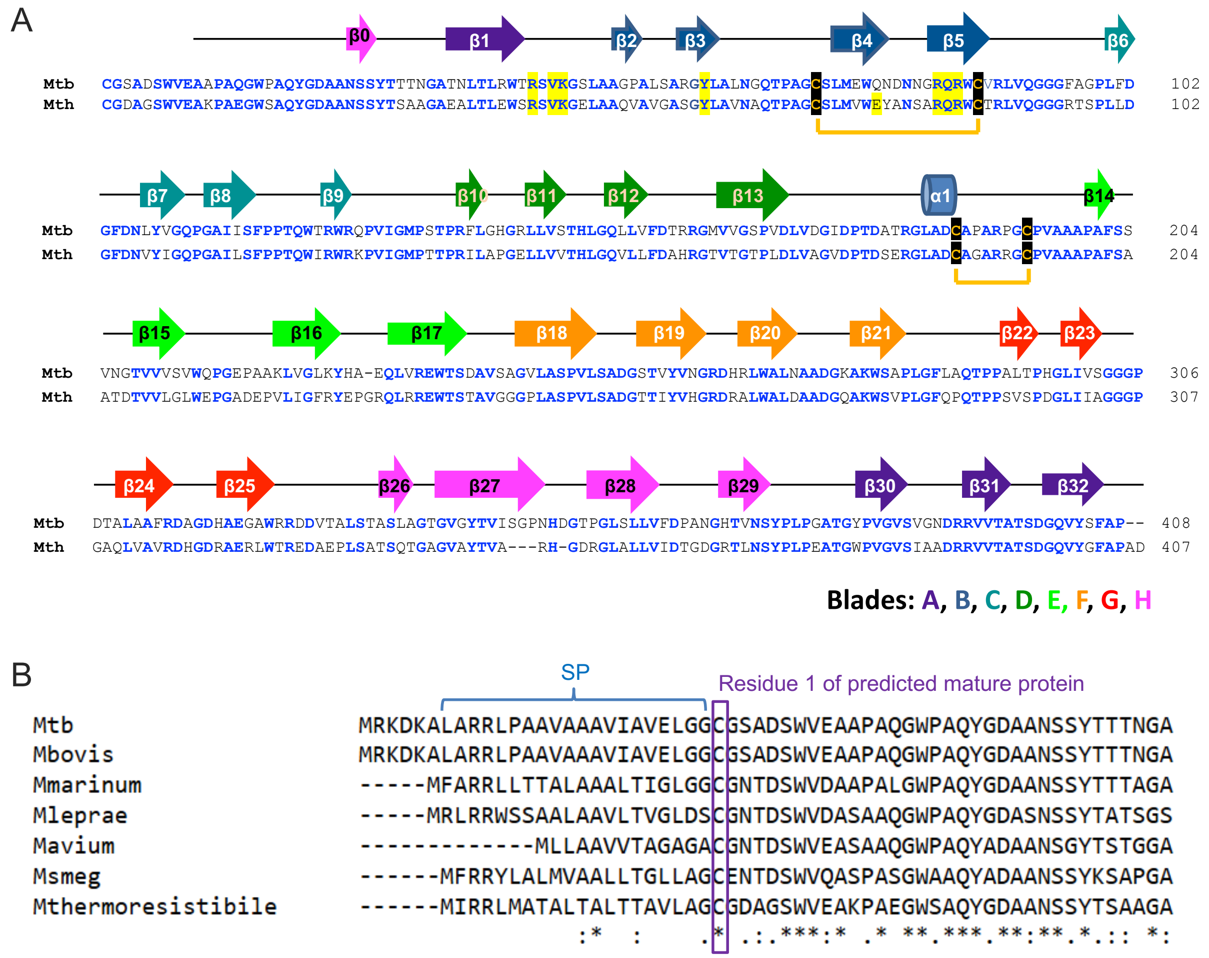


**Fig. S4. Rv3035 sequence alignments.** (A) Sequence alignment of Mtb and Mth Rv3035 with the secondary structure elements for the Mth Rv3035 structure shown above. The two amino acid sequences are 67% identical. β-strands are color coded to show which of the 8 blades of the β-propeller fold they form. Cys residues that participate in a disulfide bond are shown in dark yellow/black and residues that interact with FecB residues are highlighted in yellow. (B) Sequence alignment of the N-terminal amino acids of Rv3035 homologs from various mycobacterial species. As the N-termini are annotated at different locations within and between species, our Rv3035 numbering begins at the cysteine residue (in purple box) predicted to be the first residue of the mature protein following cleavage of the predicted signal peptide (SP) (SignalP (*14*)). Accession IDs: *Mycobacterium tuberculosis* (Mtb), WP_031647618; *M. bovis* (Mbovis), AMO11389; *M. marinum* (Mmarinum), RFZ02589; *M. leprae* (Mleprae), WP_197492542; *M. avium* (Mavium), APA77057; *M. smegmatis* (Msmeg), ABK70936; *M. thermoresistibile* (Mth), WP_040546537. Note that because different annotations of Rv3035 homologs use start sites other than those listed here, it is unclear what the consensus N-terminal residue is for the pre-processed protein.

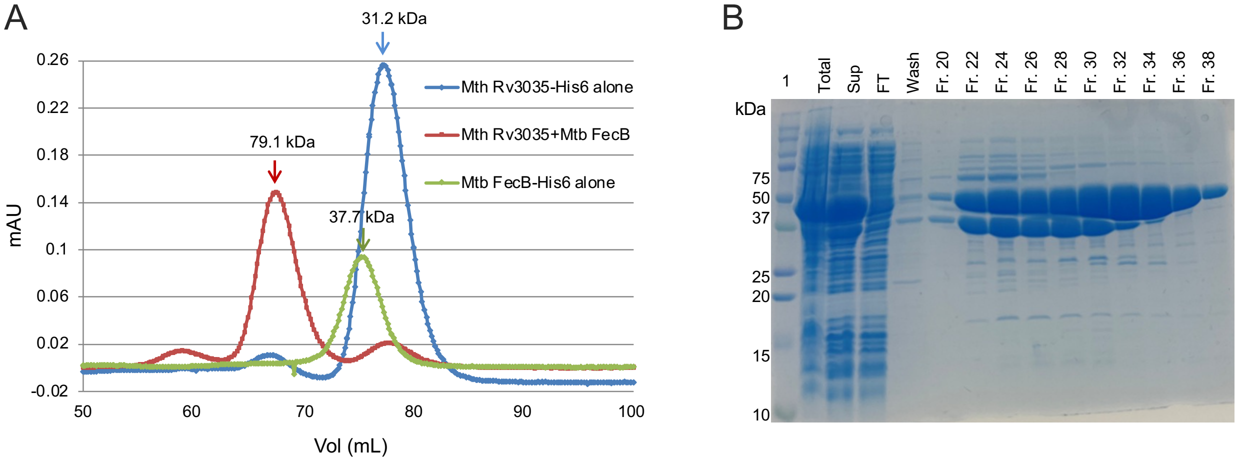


**Fig. S5. Size-exclusion chromatography of Mtb FecB and Mth Rv3035 alone and co-expressed.** (A) Size-exclusion chromatography (Superdex 200) of purified Mth Rv3035-His6 alone, Mtb FecB-His6 alone, and the Mth Rv3035-Mtb FecB complex (with Rv3035-His6 and Mtb FecB untagged). Apparent molecular weights for the indicated elution peaks were calculated using a standard curve generated from protein standards. Actual molecular weights are: Mth Rv3035-His, 44.7 kDa; Mtb FecB (untagged), 34.4 kDa; Mtb FecB-His, 36.6 kDa; Mth Rv3035 (tagged) + Mtb FecB (untagged), 79.1 kDa. Notably, Mth Rv3035 alone runs with an apparent molecular weight considerably smaller than its actual molecular weight, which may be due to the compact nature of the Rv3035 structure. PAGE analysis of the SEC fractions showed that Rv3035 runs at the correct molecular weight following SEC indicating the protein is not degraded (not shown). (B) SDS-PAGE gel of Ni-affinity purified co-expression of Mth Rv3035-His and Mtb FecB. Total = unlysed cells, Sup = supernatant minus cell debris; FT = flow-through. Elution fractions (Fr) are indicated and show that Rv3035 and FecB form a stable complex. Lane 1 is a protein ladder. kDa = kilodalton.

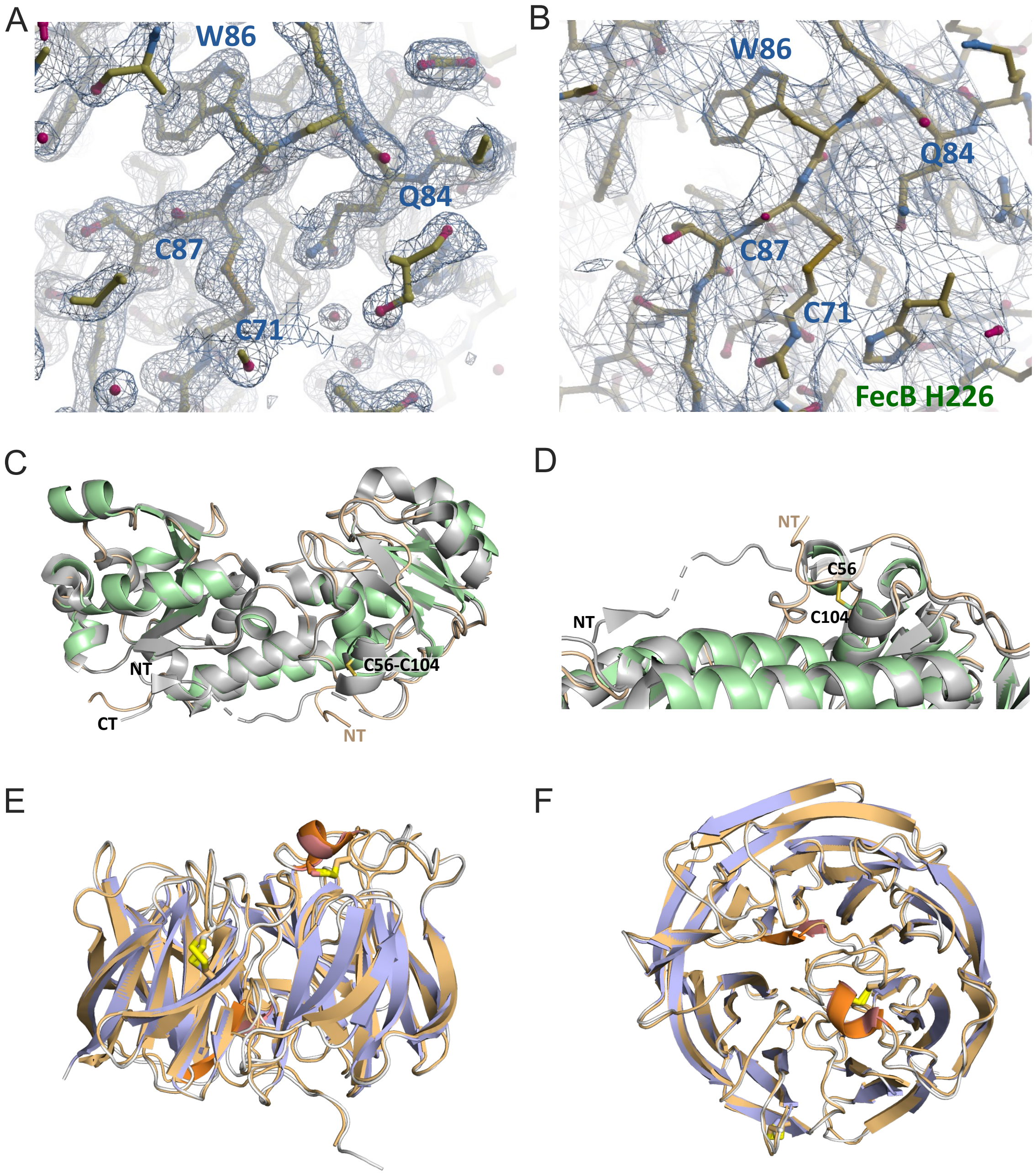


**Fig. S6. Representative electron density maps for Rv3035 alone and in complex with FecB, and superimposition of the FecB-Rv3035 complex with FecB or Rv3035 alone.** (A-B) Representative 2Fo-Fc electron density maps surrounding the C71-C87 disulfide bond contoured at $1\alpha$ for (A) Rv3035 alone (1.9 Å) and (B) the FecB-Rv3035 complex at (3.35 Å). (C-D) Structural comparison of FecB alone (gray, PDB code: 7UQ0) to FecB in complex with Rv3035 (light green). Apo-FecB aligns to FecB in complex with Rv3035 with 0.5 Å rmsd over 296 residues. (E-F) Structural comparison of Rv3035 alone (light orange and dark orange) to Rv3035 (light purple and pink) in complex with FecB. There is very little structural difference between Rv3035 and Rv3035 in complex with FecB with a 0.85 Å rmsd over 392 residues.

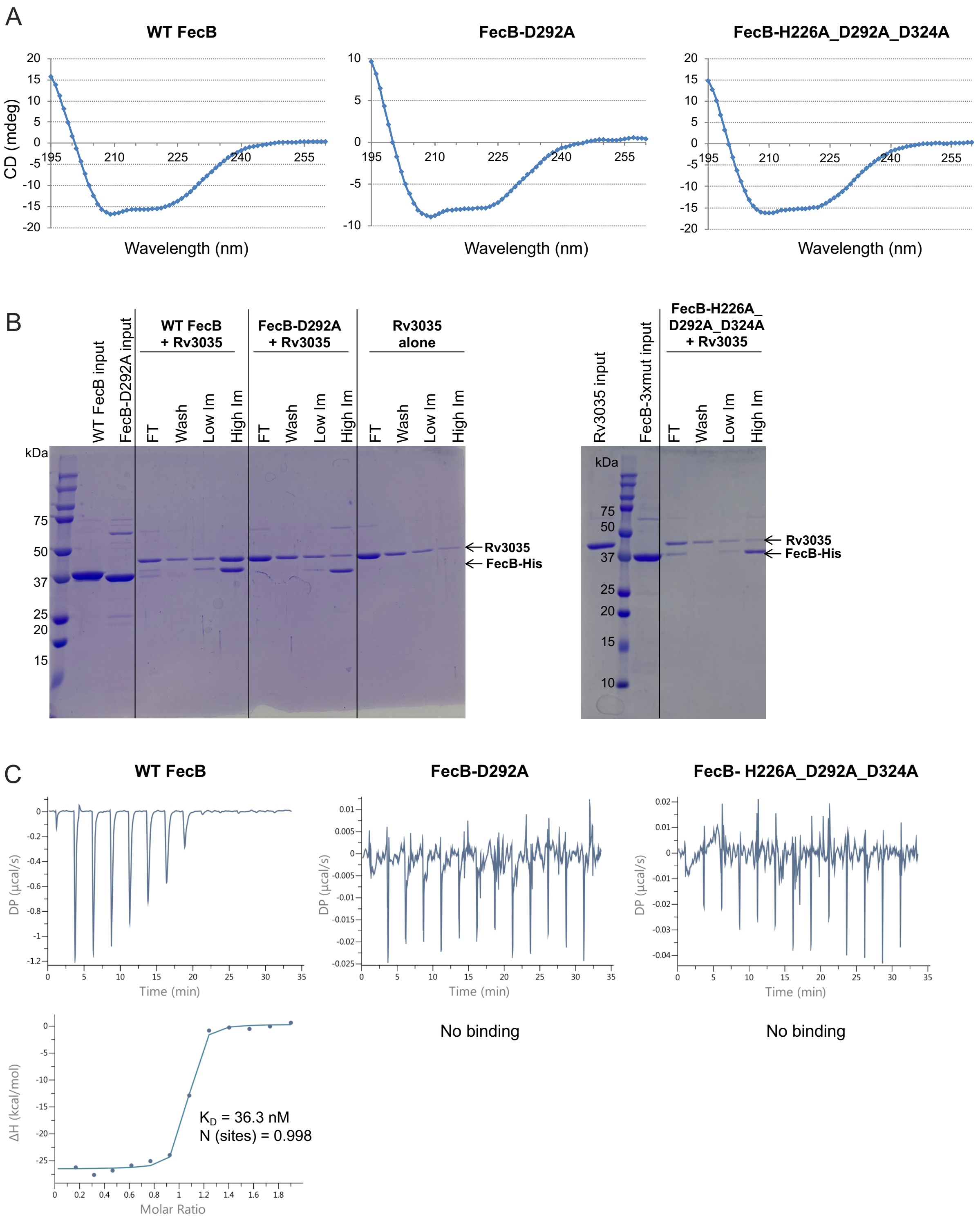


**Fig. S7. Characterization of FecB mutants D292A and H226A_D292A_D324A *in vitro*.** (A) Circular dichroism spectra of purified WT FecB, FecB D292A and FecB-H226A_D292A_D324A. All proteins have His-tags; all appear to be correctly folded. (B) WT, mutant, or no FecB-His protein was incubated with untagged Rv3035 and Ni-NTA resin. Resin was washed with binding buffer (Wash), buffer + 50 mM imidazole (Low Im), or buffer + 400 mM imidazole (High Im). Binding of Rv3035 to FecB is indicated when untagged Rv3035 is present in the high Im elution at concentrations above the background levels seen in the Rv3035 alone control. The results show that WT FecB-His and Rv3035 form a protein-protein interaction; however, the single FecB-D292A mutant and the triple mutant only pull down background levels of Rv3035, suggesting that these FecB mutants no longer bind to Rv3035. (C) Isothermal titration calorimetry (ITC) experiments using Rv3035 with WT (top, left), FecB-D292A (top, middle), or FecB-H226A_D292A_D324A (top, right) FecB-His. 13-point ITC experiments were carried out using 200μM Mth Rv3035 and 20μM Mtb WT FecB-His or 300μM Mth Rv3035 and 30μM purified FecB mutant proteins (His-tagged). The resulting thermograms are shown, indicating strong binding between WT FecB with Rv3035 and no binding of FecB-D292A or FecB-H226A_D292A_D324A with Rv3035. A binding isotherm (bottom, left) was fit to the data from WT FecB resulting in the following calculated parameters: $K_D$ = 36.3 +/- 14.2 nM; N(sites) = 0.998 +/- 0.0067.

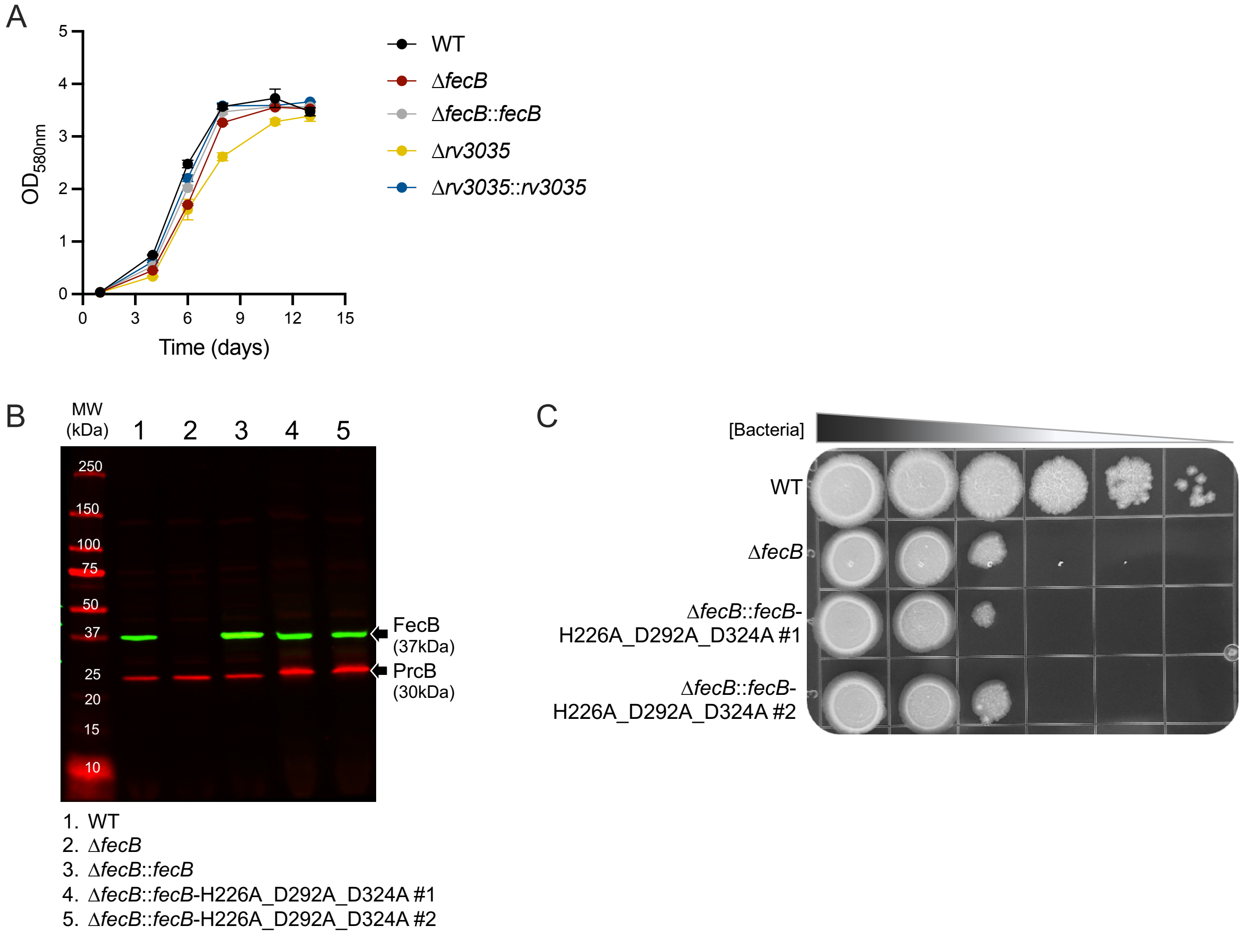


**Fig. S8. Growth curve of Δ*rv3035* and characterization of Δ*fecB* complemented with the *fecB*-H226A_D292A_D324A mutant.** Related to Fig. 5. (A) Growth curve of indicated strains using regular media in standing flasks. (B) Western-blot analysis of whole cell lysates. Nitrocellulose membrane was probed with anti-FecB antiserum and anti-PrcB primary antibody (loading control). MW = molecular weight, kDa = kilodalton. (C) Spot Assay. #1-2 after strain name indicate individual colonies (biological replicates). Error bars: ±SD.

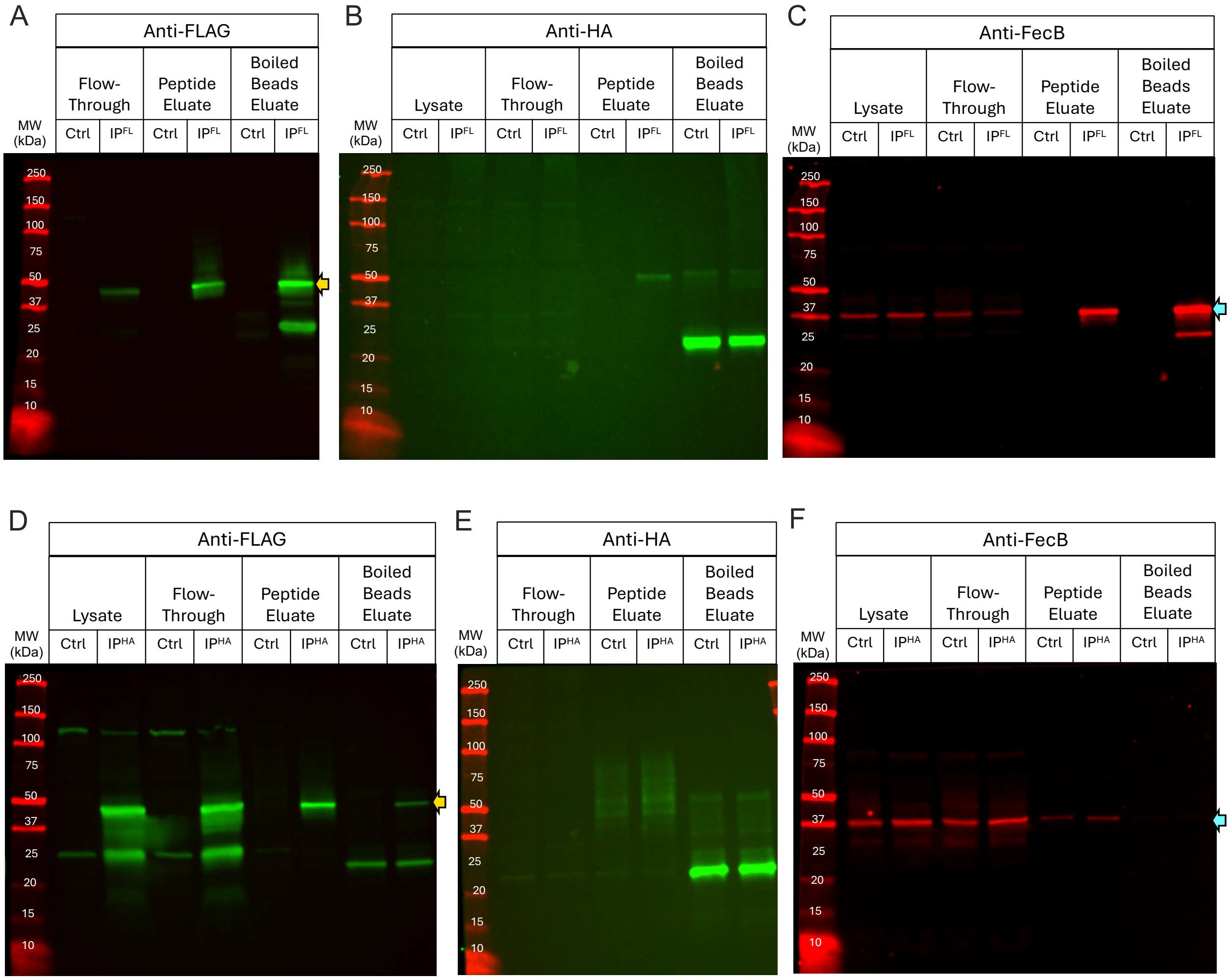


**Fig. S9. Co-immunoprecipitation of Rv3035-3xFLAG and HA-AftB.** Western blot analysis of protein fractions from Δ*rv3035* harboring plasmids encoding Rv3035-3xFLAG and HA-AftB immunoprecipitated with FLAG (A-C, $IP^{FL}$) or HA (D-F, $IP^{HA}$) magnetic beads. WT lysate without tagged proteins was used as a control (Ctrl). Nitrocellulose membranes were probed with anti-FLAG primary antibody (A, D) to detect Rv3035 (~44kDa, yellow arrows), anti-HA primary antibody (B, E) to detect AftB (~69kDa, not visualized), and anti-FecB antiserum (C, F) to detect FecB (~37kDa, blue arrows). MW = Molecular Weight, kDa = kilodalton.

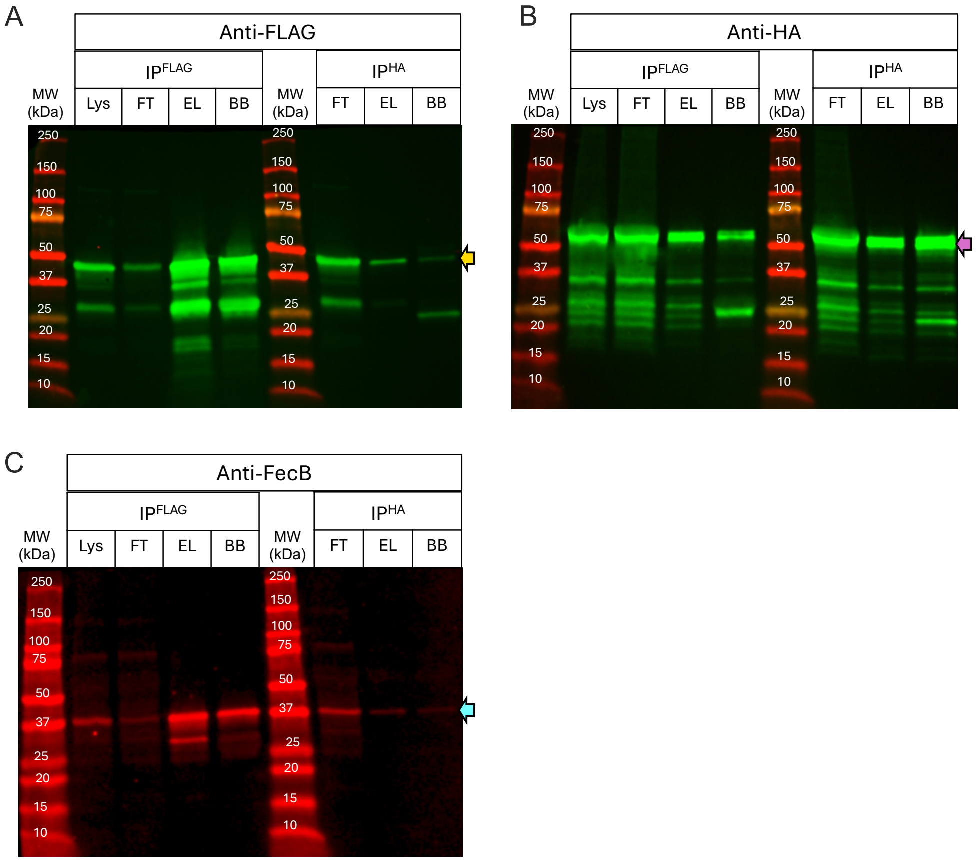


**Fig. S10. Co-immunoprecipitation of Rv3035-3xFLAG and Rv0227c-HA.** Western blot analysis of protein fractions from Δ*rv3035* harboring plasmids encoding Rv3035-3xFLAG and Rv0227c-HA immunoprecipitated with FLAG ($IP^{FLAG}$) or HA ($IP^{HA}$) magnetic beads. Nitrocellulose membranes were probed with anti-FLAG primary antibody (A) to detect Rv3035 (~44kDa, yellow arrow), anti-HA primary antibody (B) to detect Rv0227c (~46kDa, observed at higher molecular weight, purple arrow), and anti-FecB antiserum (C) to detect FecB (~37kDa, blue arrow). MW = Molecular Weight, kDa = kilodalton, Lys = lysate, FT = flow-through, EL = peptide eluate, BB = boiled beads eluate.

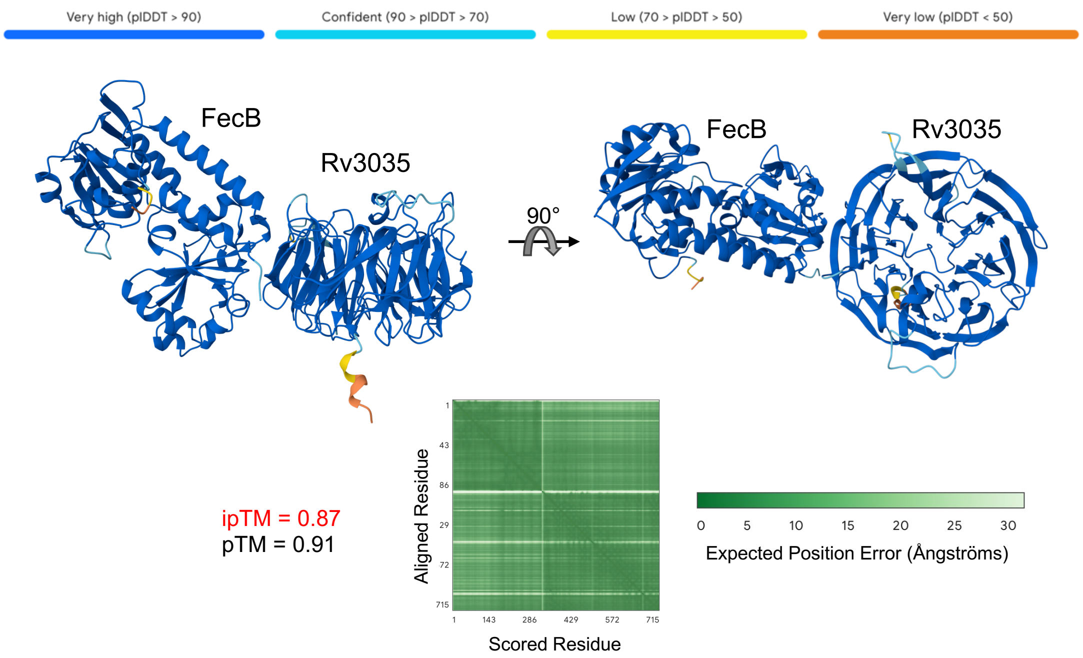


**Fig. S11. AlphaFold model of the FecB-Rv3035 complex.** AlphaFold 3 (*26*) protein complex modeling of FecB and Rv3035 from Mtb. Colored bars at the top represent pLDDT per-atom confidence estimate (dark blue indicates highest confidence). Graphs represent predicted aligned error (darker green represents lower predicted error and therefore higher confidence). Interface predicted template modeling (ipTM) scores higher than 0.8 (highlighted in red) indicate confident high-quality predictions, while values below 0.6 indicate failed predictions. Predicted template modeling (pTM) score above 0.5 indicates similarity of the predicted complex fold to the true structure.

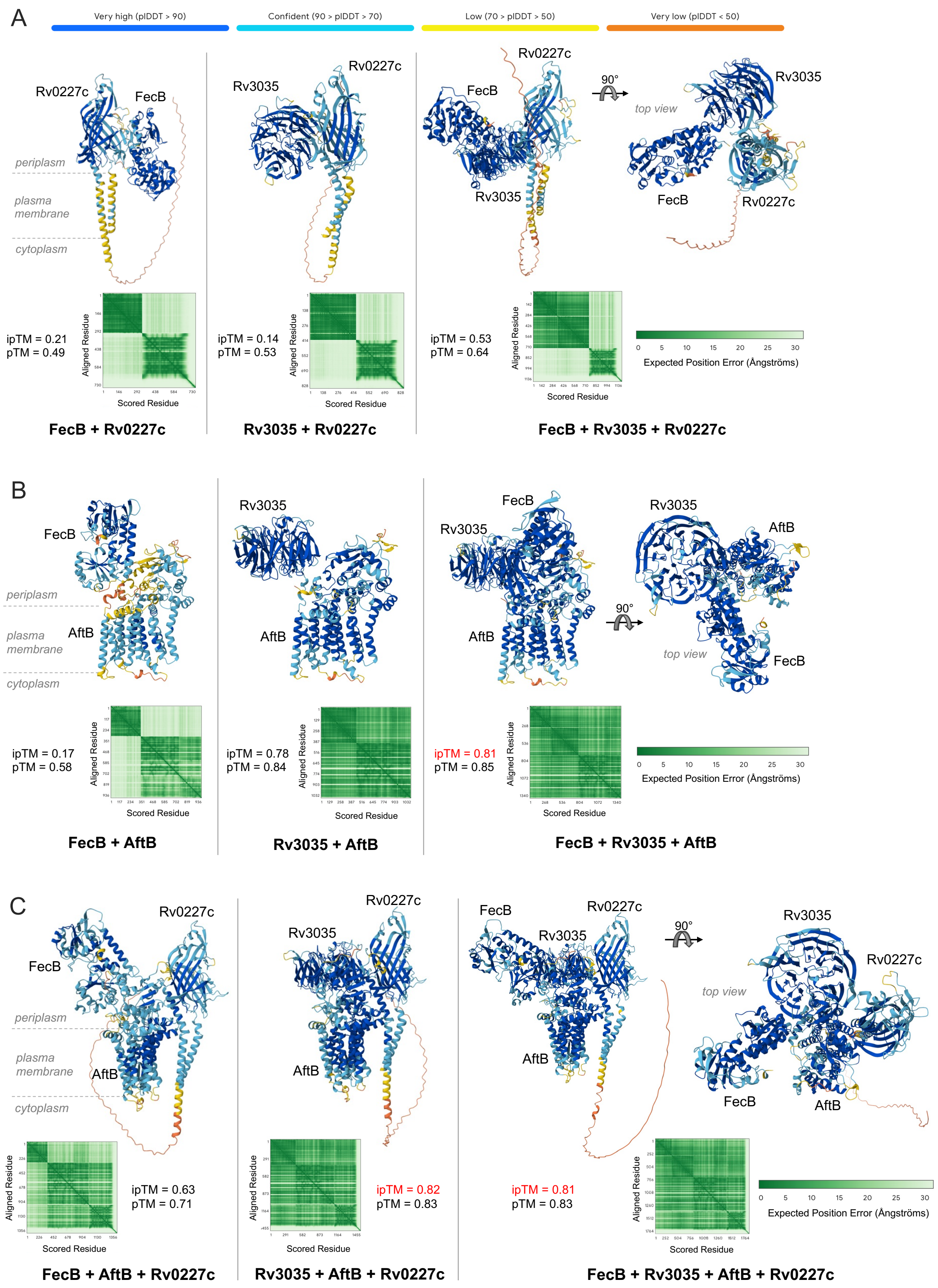

A
Very high (pIDDT > 90)
Confident (90 > pIDDT > 70)
Low (70 > pIDDT > 50)
Very low (pIDDT < 50)
Rv0227c
FecB
periplasm
plasma membrane
cytoplasm
ipTM = 0.21
pTM = 0.49
FecB + Rv0227c
Rv3035
Rv0227c
ipTM = 0.14
pTM = 0.53
Rv3035 + Rv0227c
FecB
Rv0227c
Rv3035
90°
top view
ipTM = 0.53
pTM = 0.64
Expected Position Error (Ångströms)
FecB + Rv3035 + Rv0227c
Aligned Residue
Scored Residue
B
FecB
AftB
ipTM = 0.17
pTM = 0.58
FecB + AftB
Rv3035
AftB
ipTM = 0.78
pTM = 0.84
Rv3035 + AftB
ipTM = 0.81
pTM = 0.85
FecB + Rv3035 + AftB
C
ipTM = 0.63
pTM = 0.71
FecB + AftB + Rv0227c
ipTM = 0.82
pTM = 0.83
Rv3035 + AftB + Rv0227c
ipTM = 0.81
pTM = 0.83
FecB + Rv3035 + AftB + Rv0227c

**Fig. S12. AlphaFold models of FecB, Rv3035, AftB and Rv0227c multimers.** AlphaFold 3 (*26*) protein complex modeling of (A) FecB and Rv3035 with Rv0227c alone or in combination, (B) FecB and Rv3035 with AftB alone or in combination and (C) FecB and Rv3035 with AftB and Rv0227c alone or in combination. Colored bars at the top represent pLDDT per-atom confidence estimate (dark blue indicates highest confidence). Graphs represent predicted aligned error (darker green represents lower predicted error and therefore higher confidence). Interface predicted template modeling (ipTM) scores higher than 0.8 (highlighted in red) indicate confident high-quality predictions, while values below 0.6 indicate failed predictions. Predicted template modeling (pTM) score above 0.5 indicates similarity of the predicted complex fold to the true structure.

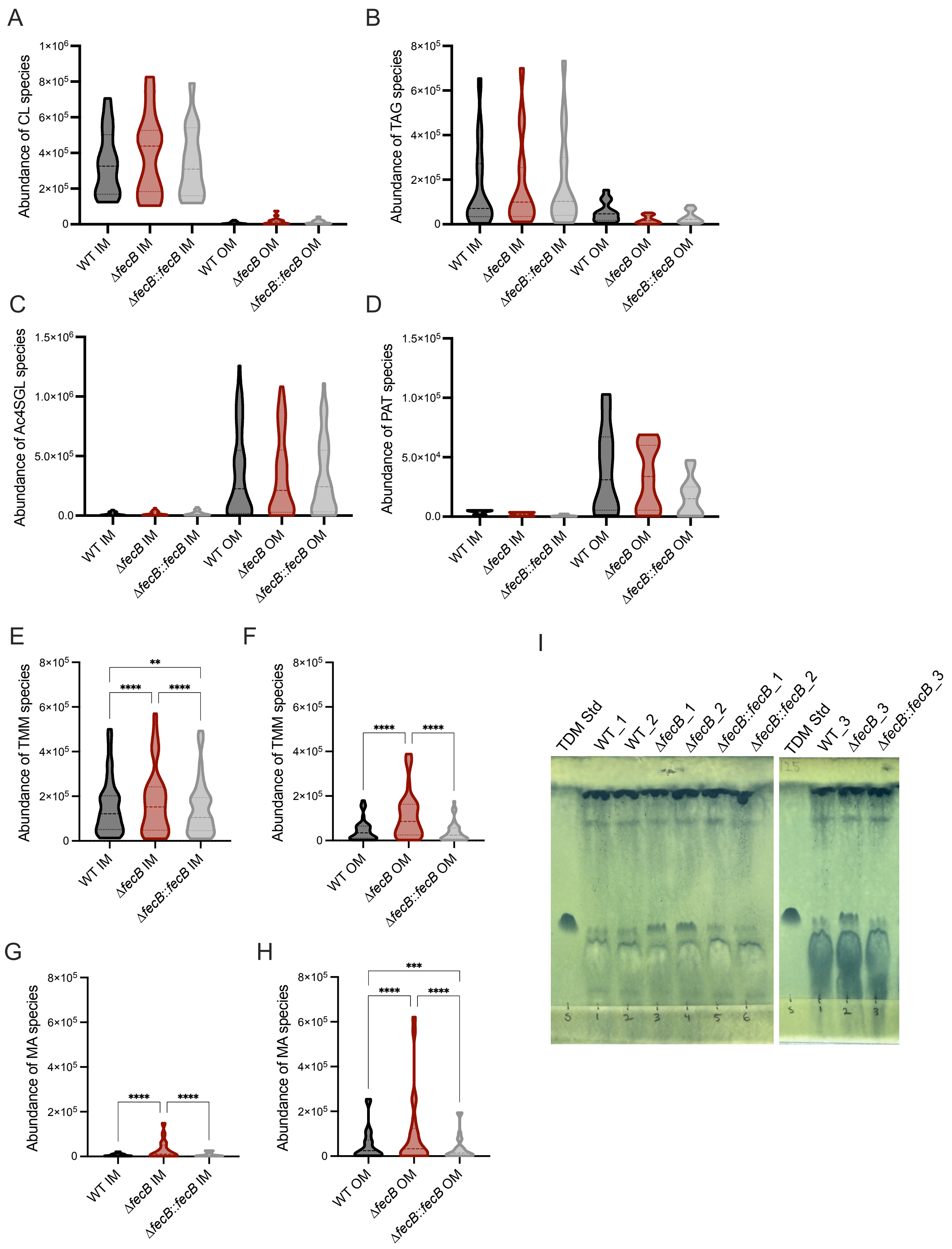
A
Abundance of CL species
1×10^6
8×10^5
6×10^5
4×10^5
2×10^5
0
WT IM
ΔfecB IM
ΔfecB::fecB IM
WT OM
ΔfecB OM
ΔfecB::fecB OM
B
Abundance of TAG species
8×10^5
6×10^5
4×10^5
2×10^5
0
WT IM
ΔfecB IM
ΔfecB::fecB IM
WT OM
ΔfecB OM
ΔfecB::fecB OM
C
Abundance of Ac4SGL species
1.5×10^6
1.0×10^6
5.0×10^5
0.0
WT IM
ΔfecB IM
ΔfecB::fecB IM
WT OM
ΔfecB OM
ΔfecB::fecB OM
D
Abundance of PAT species
1.5×10^5
1.0×10^5
5.0×10^4
0.0
WT IM
ΔfecB IM
ΔfecB::fecB IM
WT OM
ΔfecB OM
ΔfecB::fecB OM
E
Abundance of TMM species
8×10^5
6×10^5
4×10^5
2×10^5
0
**
****
****
WT IM
ΔfecB IM
ΔfecB::fecB IM
F
Abundance of TMM species
8×10^5
6×10^5
4×10^5
2×10^5
0
****
****
WT OM
ΔfecB OM
ΔfecB::fecB OM
G
Abundance of MA species
8×10^5
6×10^5
4×10^5
2×10^5
0
****
****
WT IM
ΔfecB IM
ΔfecB::fecB IM
H
Abundance of MA species
8×10^5
6×10^5
4×10^5
2×10^5
0
***
****
****
WT OM
ΔfecB OM
ΔfecB::fecB OM
I
TDM Std
WT_1
WT_2
ΔfecB_1
ΔfecB_2
ΔfecB::fecB_1
ΔfecB::fecB_2
TDM Std
WT_3
ΔfecB_3
ΔfecB::fecB_3

**Fig. S13. Lipid analysis of ∆*fecB* and control strains after AOT treatment.** Lipidomics analysis by LC-MS of fractions from AOT treatment. (A-D) Validation of representative lipids enriched in the IM (A-B) and OM (C-D) fractions. (A) cardiolipin (CL), (B) triacylglycerol (TAG), (C) sulfolipid (Ac4SGL), (D) pentaacyl trehalose (PAT). (E-F) Trehalose monomycolate (TMM) species in (E) IM and (F) OM fractions. (G-H) free-mycolic acid (MA) species in (G) IM and (H) OM fractions. (I) Thin-layer chromatography plate for detection of TDMs using AOT treatment OM fractions of WT, ∆*fecB* and ∆*fecB*::*fecB* strains. TDM standard (Std) was used to identify the TDM band location for densitometry analysis. Statistical significance was determined by two-way ANOVA and Tukey post-hoc test. N independent experiments = 1, performed with triplicate cultures. Error bars: ±SD. *$P$ < 0.05; **$P$ < 0.01; ***$P$ < 0.001; ****$P$ <0.0001.

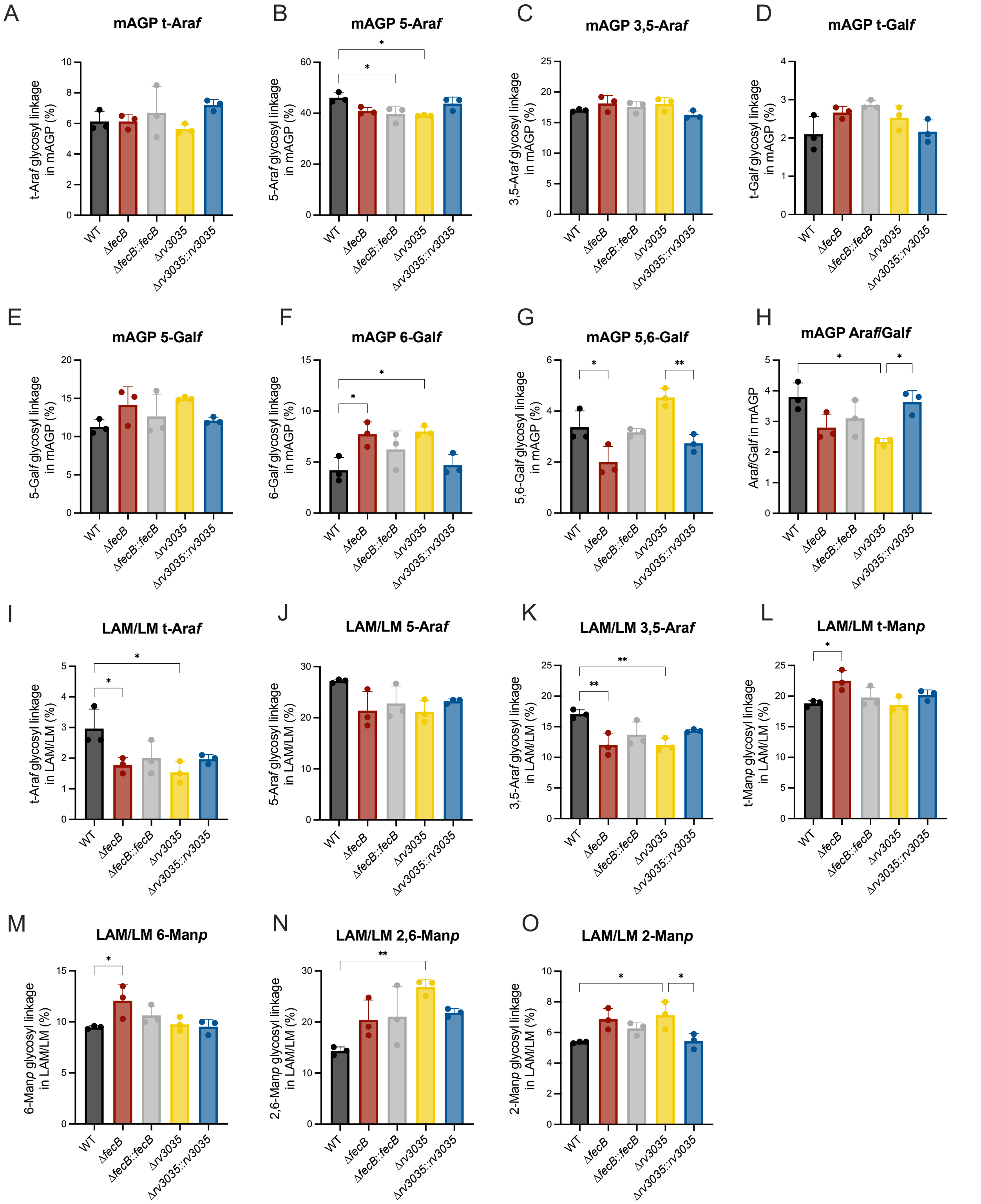


**Fig. S14. Glycosyl linkage analysis of per-*O*-methylated mAGP, LAM and LM.** The Y-axis indicates the relative percentage of each type of glycosyl residue in the (A-H) mAGP complex and (I-O) lipoglycans (LAM and LM mixture) from WT, Δ*fecB*, Δ*fecB*::*fecB*, Δ*rv3035* and

Δ*rv3035*::*rv3035* strains. The values for 2-Ara*f* residues are presented in Fig. 6, H and I. Statistical significance was determined by one-way ANOVA and Tukey post-hoc test. N independent experiments = 1, performed with triplicate cultures. Error bars: ±SD. *$P$ < 0.05; **$P$ < 0.01; ***$P$ < 0.001; ****$P$ <0.0001.

### Supplementary tables

**Table S1. Data collection and refinement statistics for Mth Rv3035 alone and the FecB-Rv3035 complex.**

| | Mth Rv3035 | FecB-Rv3035 |
|---|---|---|
| **Data collection** | | |
| Space group | $P\ 4_3\ 2_1\ 2$ | $P\ 3_2\ 2\ 1$ |
| Cell dimensions | | |
| *a*, *b*, *c* (Å) | 69.7, 69.7, 327.9 | 154.9, 154.9, 98.9 |
| α, β, γ (°) | 90, 90, 90 | 90, 90, 120 |
| Wavelength (Å) | 1.0 | 1.0 |
| Resolution (Å)[a] | 39.40-1.90 (1.94-1.90) | 39.80-3.35 (3.62-3.35) |
| $R_{merge}$[b] | 0.192 (3.020) | 0.258 (2.724) |
| $CC_{1/2}$ | 0.999 (0.867) | 0.998 (0.806) |
| $I / \sigma I$ | 11.7 (1.8) | 7.9 (1.6) |
| Completeness (%) | 100 (100) | 99.9 (100.0) |
| Redundancy | 18.5 (18.8) | 22.7 (22.0) |
| **Refinement** | | |
| Resolution (Å) | 39.40-1.90 (1.92-1.90) | 39.80-3.35 (3.47-3.35) |
| No. reflections | 64964 (2099) | 39916 (3878) |
| $R_{work}$ / $R_{free}$[c] | 17.6/21.4 | 22.8/25.4 |
| Ramachandran favored (%) | 96.85 | 95.44 |
| Ramachandran outliers (%) | 0 | 0 |
| No. atoms | | |
| Protein | 5846 | 4712 |
| Ligand/ion | 107 | 19 |
| Water | 432 | 18 |
| *B*-factors | | |
| Protein | 33.43 | 142.65 |
| Ligand/ion | 45.9 | 137.95 |
| Water | 38.59 | 111.90 |
| R.m.s deviations | | |
| Bond lengths (Å) | 0.006 | 0.006 |
| Bond angles (°) | 0.86 | 0.82 |
| PDB identifier | 9P3G | 9P3F |

[a] Values within parentheses refer to the highest resolution shell.

[b] $R_{merge} = \sum\sum|I_{hkl} - I_{hkl}(j)|/\sum I_{hkl}$, where $I_{hkl}(j)$ is observed intensity and $I_{hkl}$ is the final average value of intensity.

[c] $R_{work} = \sum||F_{obs}| - |F_{calc}||/\sum|F_{obs}|$ and $R_{free} = \sum||F_{obs}| - |F_{calc}||/\sum|F_{obs}|$, where all reflections belong to a test set of 10% data randomly selected in Phenix.

**Table S2. Structural homologs for Mth Rv3035 using Dali** (*68*)

| **Structural homolog** | **Identity (%)** | **PDB ID** | **Z-score** | **rmsd, Å** | **Ref.** |
|---|---|---|---|---|---|
| Plant growth-promoting factor YxaL from *Bacillus velezensis* | 18 | 7EQ5 | 39.4 | 2.3 (339 residues) | (*69*) |
| Dynein assembly transport adaptor from *Chlamydomonas reinhardtii* | 10 | 5MZH | 34.0 | 2.4 (324 residues) | (*70*) |
| Tetrathionate hydrolase from *Acidithiobacillus ferrooxidans* | 15 | 7CQY | 32.2 | 2.4 (328 residues) | (*71*) |
| Rsa4 from *Chaetomium thermophilum* | 11 | 4WJS | 32.2 | 2.5 (325 residues) | (*72*) |
| Outer membrane assembly protein BamB in BAM-SurA complex from *E. coli* | 20 | 9HG9 | 31.8 | 2.8 (317 residues) | (*73*) |
| BamB-BamA PORTA domain from *E. coli* | 20 | 4XGA | 31.7 | 2.8 (315 residues) | (*74*) |
| Methanol Dehydrogenase from *Methylacidiphilum fumariolicum* | 18 | 6FKW | 31.5 | 3.1 (354 residues) | (*75*) |
| SQT1 chaperone of ribosome protein to protect the rRNA binding residues from *Thermochaetoides thermophila* | 13 | 4ZN4 | 30.2 | 2.6 (339 residues) | (*76*) |
| Dos1 propeller from *Schizosaccharomyces pombe* | 11 | 4O9D | 29.8 | 3.2 (334 residues) | (*77*) |
| PQQ-dependent alcohol dehydrogenase detoxifying DON from *Devosia albogilva* | 14 | 7WMD | 29.3 | 3.0 (354 residues) | (*78*) |

**Table S3. Interaction distances for hydrogen bonds and salt bridges Rv3035 and FecB with interactions $\leq$ 3.5 Å.** Interactions that are < 3.0 Å highlighted in blue.

| **Rv3035** | | | **FecB** | | | Distance (Å) |
|---|---|---|---|---|---|---|
| Res Number | Res Name | Atom | Res Number | Res Name | Atom | |
| 43 | Arg | NH1 | 292 | Asp | Oδ1 | 3.5 |
| 43 | Arg | NH1 | 292 | Asp | Oδ2 | 3.5 |
| **43** | **Arg** | **NH2** | **292** | **Asp** | **Oδ2** | **2.7** |
| 45 | Val | O | 226 | His | Nε2 | 3.2 |
| 46 | Lys | O | 226 | His | Nε2 | 3.5 |
| 46 | Lys | Nζ | 258 | Asp | Oδ2 | 3.1 |
| 60 | Tyr | OH | 323 | Arg | NH2 | 3.4 |
| 71 | Cys | N | 223 | Asp | Oδ2 | 3.3 |
| 77 | Glu | Oε1 | 320 | Ser | Oγ | 3.3 |
| **77** | **Glu** | **Oε2** | **323** | **Arg** | **NH2** | **2.8** |
| **83** | **Arg** | **NH1** | **291** | **Ala** | **O** | **2.8** |
| 83 | Arg | NH1 | 292 | Asp | O | 3.4 |
| 83 | Arg | NH2 | 320 | Ser | O | 3.4 |
| **83** | **Arg** | **NH2** | **324** | **Asp** | **Oδ2** | **2.8** |
| **84** | **Gln** | **N** | **292** | **Asp** | **Oδ1** | **2.9** |
| **84** | **Gln** | **Nε2** | **226** | **His** | **O** | **2.7** |
| 85 | Arg | NH2 | 324 | Asp | Oδ1 | 3.3 |
| 85 | Arg | Nε | 324 | Asp | Oδ1 | 3.3 |
| 85 | Arg | Nε | 324 | Asp | Oδ2 | 3.3 |

**Table S4. Primers and probes used for RT qPCR**

| Gene name and Rv# | ID | Type of sequence | Sequence | Labeling |
|---|---|---|---|---|
| *mbtB* (Rv2383c) | mbtB-fw | Forward primer | CCGGGAGATGTTGACCAGTTG | |
| | mbtB-rv | Reverse primer | GCGTCGAGGTGATCGTTGAGAT | |
| | mbtB-prb | Probe | AGACTCACACCTCCGGTCGCTTC | 5' FAM / 3' BBQ-1 |
| *ppe37* (Rv2123) | ppe37_fw | Forward primer | TGTCCGGTCCAGTCTTCAC | |
| | ppe37_rv | Reverse primer | GCGAACGGTCCGAAATAGATCAG | |
| | ppe37_prb | Probe | TTTCTCGCCTACCTGGTGCTGG | 5' FAM / 3' BHQ1 |
| *irtA* (Rv1348) | irtA-fw | Forward primer | AAACCTGGCGCAACCATA | |
| | irtA-rv | Reverse primer | GAGTCGCCGATTAGCAGATAC | |
| | irtA-prb | Probe | TGAGCCCATCAGCGACATGACC | 5' FAM / 3' BHQ-1 |
| *fecB* (Rv3044) | fecb-fw | Forward primer | CGAACAGCTCCTCAAGTCAA | |
| | fecb-rv | Reverse primer | CTGCGAACCCAGGATCAG | |
| | fecb-prb | Probe | ATGATCTGCCCGGTGTCGGTAC | 5' 6FAM / 3' BBQ |

**Table S5. Primers used to produce FecB mutants**

| Name | Sequence |
|---|---|
| FecB-H226A-forward | CATGACGCGACCGCCTTCCAAGCGTCG |
| FecB-H226A-reverse | CGACGCTTGGAAGGCGGTCGCGTCATG |
| FecB-D292A-forward | CCGACGCCGCTATCGTCTACCTGTC |
| FecB-D292A-reverse | GACAGGTAGACGATAGCGGCGTCGG |
| FecB-D324A-forward | GCCAACCGTGCCAACCGGGTCTTCG |
| FecB-D324A-reverse | CGAAGACCCGGTTGGCACGGTTGGC |

**Table S6. Primers used to confirm allelic exchange in Δ*rv3035* and Δ*fecB*Δ*rv3035* strains**

| Name | Sequence |
|---|---|
| Rv3035-UHR_Fw | TCGGACCTGTTTGGGCG |
| Rv3035-DHR_Rv | TGACCGACTACATCACGCAAAAC |
| seq-Zeo-F1 | AAGTTGACCAGTGCCGTTCC |
| seq-Zeo-R1 | GCGCTGATGAACAGGGTCAC |
| Rv3035-UHR_Fw2 | GATCCTCGCCACCTGCGG |
| Rv3035-DHR_Rv2 | GTTGCTGTGGTCCTGGTGG |

**Supplementary datasets**

**Data S1 (separate file).** Related to Fig 1. RNA-seq normalized counts and differential expression analysis results for WT, Δ*fecB* and Δ*fecB*::*fecB* strains grown in detergent-free (DF) or Tween80-containing media.

**Data S2 (separate file).** Related to Fig 3. Total spectrum count analysis of pull-down tandem mass spectrometry experiment for FecB-FLAG and Rv3035-3xFLAG, including overlapping hits determined as proteins with >= 1 log2FC relative to the WT lysate control.

**Data S3 (separate file).** Related to Fig 3. Co-essentiality analysis results for query genes *fecB*, *rv3035* and *aftB*.

**Data S4 (separate file).** Related to Fig 6. Targeted lipid analysis of AOT-treated (OM and IM fractions) and untreated samples.

**Data S5 (separate file).** Related to Material and Methods. List of Mtb strains used in this study. Plasmids are shown in full strain name column after the double collon. For plasmids constructed using the Invitrogen Gateway Cloning method, pGM refers to the destination vector pDE43-M; after that, C/E indicates the type of plasmid, with C for chromosomally integrated (the absence of a lower case letter after C refers to integration at attL5, while t indicates integration at the tweety site) or E for episomal; selective antibiotic resistance conferred by the plasmid is indicated by the last letter before the dash (e.g. pGMCK, for kanamycin). Resistance conferred by each plasmid: Hyg/H, hygromycin; Kan/K, kanamycin; Strep/S, streptomycin; Zeo/Z, zeocin; Apra/A, apramycin. Both T02 and OXOX refer to an empty insert (*79*). XSTS refers to a plasmid that contains colorimetric selection element used in selection, *xylE*, as well as a gene for counterselection on sucrose, *sacB* (*80*, *81*). Phsp60 is a constitutive promoter. T10M refers to a TetR expression cassette and P606.10C is a tetO-containing promoter of intermediate strength. T10M mediates repression of P606.10C, which is relieved by anhydrotetracycline. The Shine-Dalgarno (SD) sequence in plasmids containing SD1 is AGGAGG. pIRL58 (Addgene #166886) refers to the backbone plasmid of the CRISPRi constructs; additional details on generation of these have been previously described (*19*). Expression type refers to the regulation of the gene in the plasmid; ATc, anhydrotetracycline. sgRNA targeting sequences are shown for CRISPRi strains. In strains complemented with plasmids containing “rv3035_A”, the "A" refers to a complementation sequence that contains silent mutations rendering it resistant to targeting by "rv3035_A-sgRNA". Rv3035’s sequence in “rv3035_A-HA” encodes a protein of 360 amino acids (truncated Rv3035), while “rv3035_A-3xFLAG” encodes a protein of 434 amino acids (adjusted transcription start site), not counting the tag peptides.
